# Arabidopsis S2Lb links AtCOMPASS-like and SDG2 activity in histone H3 Lys-4 trimethylation independently from histone H2B monoubiquitination

**DOI:** 10.1101/583724

**Authors:** Anne-Sophie Fiorucci, Clara Bourbousse, Lorenzo Concia, Martin Rougée, Anne-Flore Deton-Cabanillas, Gérald Zabulon, Elodie Layat, David Latrasse, SoonKap Kim, Nicole Chaumont, Bérangère Lombard, David Stroebel, Sophie Lemoine, Ammara Mohammad, Corinne Blugeon, Damarys Loew, Christophe Bailly, Chris Bowler, Moussa Benhamed, Fredy Barneche

## Abstract

The functional determinants of histone H3 Lys-4 trimethylation (H3K4me3), their potential dependency on histone H2B monoubiquitination (H2Bub) and their contribution to defining transcriptional regimes are poorly defined in plant systems. Unlike in *S. cerevisiae*, where a single SET1 protein catalyzes H3 Lys-4 trimethylation as part of COMPASS (COMPlex of proteins ASsociated with Set1), in *Arabidopsis thaliana* this activity involves multiple histone methyltransferases (HMTs). Among these, the plant-specific SDG2 (SET DOMAIN GROUP2) has a prominent role. We report that SDG2 co-regulates hundreds of genes with SWD2-like b (S2Lb), a plant ortholog of the Swd2 axillary subunit of yeast COMPASS. S2Lb co-purifies with the AtCOMPASS core subunit WDR5 from a high-molecular weight complex, and both S2Lb and SDG2 directly influence H3K4me3 enrichment over highly transcribed genes. *S2Lb* knockout triggers pleiotropic developmental phenotypes at the vegetative and reproductive stages, including reduced fertility and seed dormancy. Notwithstanding, *s2lb* seedlings display little transcriptomic defects as compared to the large repertoire of genes targeted by S2Lb, SDG2 or H3 Lys-4 trimethylation, suggesting that H3K4me3 enrichment is important for optimal gene induction during cellular transitions rather than for determining on/off transcriptional status. Moreover, unlike in budding yeast, most of the S2Lb and H3K4me3 genomic distribution does not rely on a trans-histone crosstalk with histone H2B monoubiquitination. Collectively, this study unveils that the evolutionarily conserved COMPASS-like complex has been coopted by the plant-specific SDG2 HMT and mediates H3K4me3 deposition through an H2Bub-independent pathway in Arabidopsis.

## Introduction

Dynamic changes in chromatin organization and composition rely on many activities, such as the remodeling of nucleosome positioning, the incorporation of histone variants, DNA methylation and histone post-translational modifications (PTMs) [1–4]. Genome-wide profiling of histone PTMs in the *Arabidopsis thaliana* plant species has established that transcriptionally active genes are typically marked by acetylated histone H3 and H4, monoubiquitinated histone H2B (H2Bub) and di/trimethylated histone H3 at lysine residues such as Lys-4 and Lys-36 (H3K4me2/3, H3K36me2/3) [5–7]. Combinations of histone modifications contribute to create a multilayered system of chromatin states and transcriptional activity [8]. In plant systems a molecular mechanisms underpinning functional interdependencies between different histone modifications have recently been reported for *Polycomb*-mediated gene repression [9] but has not been characterized for active transcription.

One of the best-described cases of functional *trans*-histone crosstalk is that of H2Bub promotion of histone H3K4me3 deposition on actively transcribed genes in yeast [10] and metazoans [11]. In *S. cerevisiae* H3K4me3 deposition is catalyzed by the SET1 histone methyltransferase (HMT) embedded in a so-called COMPASS (COMPlex of Proteins Associated with Set1), which also contains the WD40-repeats-containing proteins Swd1, Swd2, Swd3 as well as Bre2, Spp1 and Sdc1 subunits (reviewed in [12, 13]). Tethering of Swd2 on H2Bub-modified nucleosomes is proposed to recruit yCOMPASS, PAF1c (Polymerase Associated Factor 1 complex) and RNA polymerase II (RNPII) to promote histone H3 Lys-4 trimethylation [14–17]. Hence, prior monoubiquitination of H2B on Lys-123 by the Rad6/Bre1 ubiquitin conjugase and E3 ligase is a prerequisite for H3K4me3 deposition by Set1 in budding yeast [10, 18–21]. COMPASS-like H3K4me3 HMT activity is evolutionarily conserved in eukaryotes with for example Trithorax (Trx) in Drosophila and Mixed Lineage Leukemia 1 (MLL1) in human [12, 22]. H3K4me3 is usually found on a limited number of nucleosomes surrounding the transcription start site (TSS) and is functionally linked to RNPII transcriptional activation and the switch to elongation in many eukaryotes including plants [5, 23–28].

As many as 47 distinct SET-domain proteins are encoded in the Arabidopsis genome, together forming the SET Domain protein Group (SDG) [29–31]. Among them, ATX1 (ARABIDOPSIS TRITHORAX1) [32] and ATXR7 (ARABIDOPSIS TRITHORAX-RELATED7) [33, 34] appear to target highly specific genomic loci or to be cell-type specific whereas the plant-specific SET DOMAIN GROUP 2/ARABIDOPSIS TRITHORAX RELATED 3 (SDG2) HMT presumably targets a broad repertoire of genes [35, 36]. Notwithstanding, while the influence of H3 Lys-4 trimethylation on transcription activation/elongation in plants have begun to emerge [37–40], the Arabidopsis genomic loci targeted by SDG2 as well as the mechanisms determining its specificity remain undetermined. ATX1 has been shown to have high affinity for Ser5-phosphorylated RNPII, a property enabling this COMPASS-associated HMT to facilitate RNPII exit from to the promoter proximal pause region to favor transcription elongation [37, 41]. A molecular crosstalk mechanistically linking H3K4me3 and H2Bub deposition has however not been established in plants [42].

In addition to HMT diversification, Arabidopsis also possesses homologs for all known COMPASS subunits such WDR5a and WDR5b playing the role of the yeast Swd3 core component, potentially forming several COMPASS-like complexes. More generally, all structural components of the yeast COMPASS (yCOMPASS) complex subunits appear to be conserved in Arabidopsis, such as a RbBP5-LIKE (RBL), a Swd3 homolog (WDR5a/b) and a Bre2 homolog (ARABIDOPSIS Ash2 RELATIVE or ASH2R) [40, 43–45]. They contribute to flowering time control by allowing H3K4me3 deposition on the *FLC (FLOWERING-LOCUS C*) regulatory gene [43, 44] but also to drought stress tolerance [46] and endoplasmic reticulum stress response [47], presumably as a consequence of a general influence on RNPII activity during cellular transitions or in response to environmental signals. Accordingly, knocking out AtCOMPASS-like core subunit genes is lethal [43, 44], indicating a fundamental role in plant development.

Here, we first identify S2Lb as an Arabidopsis homolog of the Swd2 COMPASS-associated subunit, which acts as a key component of the H2Bub-H3K4me3 *trans*-histone crosstalk in *S. cerevisiae* [14, 17]. We report that S2Lb is a euchromatic protein that functionally associates with an AtCOMPASS-like complex and with the plant-specific SDG2 HMT to broaden H3K4me3 enrichment over most transcribing genes, especially those abundantly occupied by RNPII. Using *HUB1* loss-of-function plants in which H2Bub deposition is abolished [48, 49], we further unveil that COMPASS-like activities mediate H3-Lys4 trimethylation largely independently from histone H2B monoubiquitination in Arabidopsis.

## Results

### Two evolutionarily conserved *SWD2-LIKE* genes encode euchromatic proteins in *A. thaliana*

Phylogenetic analysis of the *S. cerevisiae* Swd2 protein sequence and putative homologs in human, Drosophila and representative plant species revealed that plant *SWD2*-like genes form a distinct clade from metazoan orthologs (Figure 1a). The presence of two or more *SWD2-LIKE* genes suggests that a gene duplication event occurred before the separation of gymnosperms and angiosperms. In Arabidopsis, *At5g14530* and *At5g66240*, designated here as *SWD2-LIKE-a* (*S2La*) and *S2Lb*, encode such predicted paralogs with high amino acids sequence similarity with yeast Swd2 (45.3% and 43.4%, respectively). These two genes were first identified for their influence on Arabidopsis flowering time as *Anthesis Promoting Factor 1* (*APRF1;* [50]) and *Ubiquitin Ligase Complex Subunit 1* ([51]), respectively. RT-qPCR analysis of whole seedlings and of different adult plant organs showed that both genes are broadly expressed, *S2Lb* being usually expressed to a much higher level than *S2La* (Additional file 2 – Figure S1). This difference is also apparent in publicly available anatomy-related transcriptomes [52], with *S2La* mRNA being mildly detected in all analyzed samples except in senescent leaves (Additional file 2 – Figure S1).

**Figure 1.**
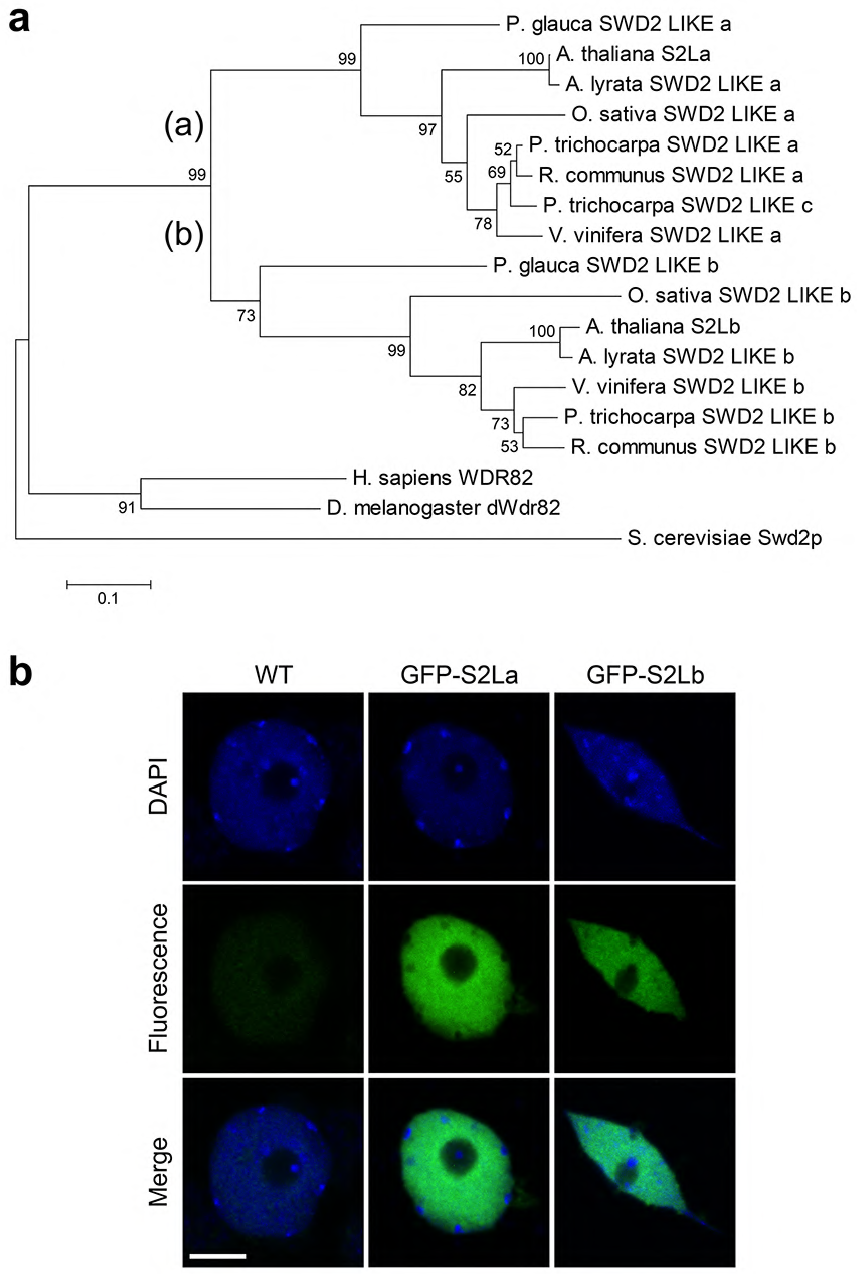
Two Swd2-like euchromatin proteins in Arabidopsis. **a** Neighbor-joining tree of full-length Swd2-Like proteins from representative species. The plant clade can be divided in two sub-clades containing each a Swd2-Like isoform, (a) and (b). Numbers indicate bootstrap values from 1,000 replicates. Scale for sequence divergence is represented by a bar. Gene and protein IDs are given in Additional file 1 (Table S1). **b** S2La and S2Lb are euchromatic proteins. Immunodetection of GFP-tagged S2La and S2Lb proteins in interphase cotyledon nuclei (green). DNA is counterstained with DAPI (blue). Scale bar, 2 µm.

In addition to this differential regulation, structural variations can be identified between S2La and S2Lb, and with their yeast and human orthologs. Six canonical WD40 repeats can be identified in S2La and S2Lb versus seven in Swd2 and in the human homolog Wdr82 (Additional file 2 – Figure S2). The WD40 repeat IV was not detected in the central part of S2La, and the S2Lb carboxy-terminal domain carries divergent WD40 repeats. Sub-nuclear immunolocalization of GFP-tagged S2La and S2Lb stably expressed *in planta* further showed that both proteins localize to euchromatin, and are excluded from the nucleolar compartment and from all densely packed heterochromatic foci known as chromocenters (Figure 1b), in agreement with a potential role linked to RNPII transcription.

### *S2Lb* but not *S2La* loss of function causes pleiotropic developmental defects

To explore the function of *S2La* or *S2Lb in planta*, T-DNA insertion lines interrupting each gene were obtained from public collections, which we named *s2la-1* [previously described as *aprf1-9* in [50]] and *s2lb-1*, respectively (Additional file 2 – Figure S3). Being in a Nossen background, the *s2lb-1* allele was introgressed in a Col-*0* background through five successive backcrosses to generate the *s2lb-2* line.

As described earlier [50], *s2la-1* plants exhibited no apparent developmental abnormalities under laboratory growth conditions at the vegetative stage. In contrast, *S2Lb* loss-of-function in both Nossen and Col-*0* backgrounds triggered significant growth defects resulting in small leaf size, small rosette diameter, shorter roots and a reduced number of lateral roots (Figure 2a-e). Such defects were not observed in heterozygous plants for the *s2lb-2* allele. Homozygous plants could efficiently be rescued by stably expressing GFP-tagged or native S2Lb proteins under control of the *S2Lb* endogenous promoter (Additional file 2 – Figure S4), thus confirming the specific and recessive property of the mutation effect on plant growth.

**Figure 2.**
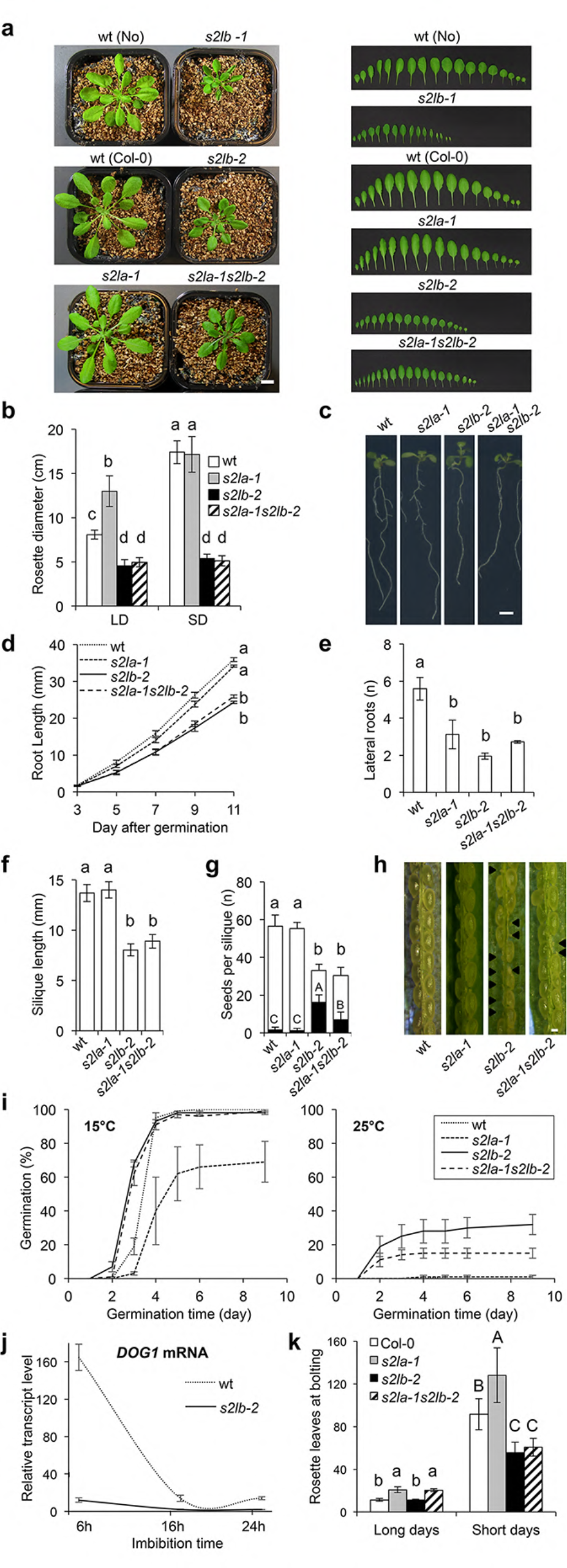
Growth defects of *s2l* mutant plants at the reproductive and vegetative stages. **a** Representative phenotypes of 4-week-old adult plants (left panel) and of dissected leaves (right panel) from plants grown under short day conditions. Scale bar, 1 cm. **b** Rosette diameter at bolting of plants grown under short (SD) or long day (LD) photoperiods. Error bars indicate SD (n>12). **c** Representative root phenotypes of 11-day-old seedlings. Scale bar, 0.5 cm. **d** Primary root length from 3 to 11 dpi. Error bars indicate SD from 2 biological replicates. **e** Seedling lateral root number at 11 dpi. Error bars indicate SD from 2 biological replicates. **f** Mean silique length and **g** number of seeds (or ovules) per silique in selfed wild-type and *s2la/b* mutant plants prior to seed desiccation. Black bars represent the mean number of undeveloped seeds per silique. Error bars indicate SD (n>12 siliques each, from 3 individuals). **h** Representative silique for each genotype. Arrows indicate undeveloped seeds/ovules. **i** Dormancy defects induced by *S2Lb* loss-of-function. The graph shows the percentage of freshly harvested germinating seeds from wild-type, *s2la*, *s2lb* and double *s2las2lb* mutant plants at 15°C (vigor test, left graph) and 25°C (dormancy test, right graph) in darkness. Error bars indicate SD between three biological replicates using two independent seed propagation sets (n=50 seeds each). **j** RT-qPCR analysis of *DOG1* transcripts level in seeds of wild-type and *s2lb* mutant plants upon imbibition. Total RNAs were extracted at the indicated time points. *DOG1* expression level relative to housekeeping genes is given as the mean of 3 independent biological replicates. **k** Flowering time phenotype analyzed as the number of leaves at bolting under SD or LD conditions. Error bars indicate SD (n>12). Letters indicate significant differences among sample means in ANOVA analysis followed by Tukey’s range test (p < 0.05).

At the reproductive stage, *s2lb-2* mutant plants display fertile flowers but the resulting siliques are short and contain a low number of ovules leading to ~50% arrested seed development (Figure 2f-h). Interestingly however, freshly harvested *s2lb-2* viable seeds reproducibly had a high germination capacity at 25°C but both genotypes displayed high seed vigor at 15°C, a temperature at which dormancy is not induced [53, 54]. These observations indicate that establishment or completion of the dormancy process is deficient in *s2lb-2* seeds (Figure 2i).

Considering the influence of the histone H2B monoubiquitination on the *DOG1* (*DELAY OF GERMINATION1*) gene [48, 55], possibly via a related chromatin mechanism, we further investigated expression of this master regulator of dormancy. As expected [56], *DOG1* was strongly expressed in imbibed seeds and subsequently downregulated during germination in wild-type plants. In contrast, *DOG1* transcripts were constantly barely detected in *s2lb-2* seeds (Figure 2j). Hence, incapacity in inducing *DOG1* upon imbibition might on its own be responsible for the dormancy phenotype.

To test for potential redundancy of *S2La* and *S2Lb* function, *s2la-1s2lb-2* double mutant plants were generated, showing largely similar vegetative phenotypic defects as *s2lb-2* single mutants (Figure 2a-e). The double mutants nonetheless generated slightly longer siliques than *s2lb-2* mutant plants (Figure 2F), producing more viable seeds (Figure 2h) with an intermediate dormancy phenotype between wild-type and *s2lb-2* seeds (Figure 2i). Strikingly, *s2la-1* and *s2lb-2* mutations also have distinct effects on flowering time control under a long-day photoperiod [as shown previously with *s2la-1* (*aprf1-9* in [50]) and *S2Lb* RNAi lines [51]]. Under a short-day photoperiod, the *s2la-1* plants exhibited a late flowering phenotype while *s2lb-2* and double mutant plants were early flowering (Figure 2k). Collectively, these analyses indicate that i) *S2Lb* is more expressed than *S2La*, ii) has a more pleiotropic role than *S2La* in vegetative and reproductive phases of plant development and iii) some of the phenotypes observed in *s2lb* mutant can be partially rescued by *S2La* loss-of-function.

### *S2Lb* is a major determinant of the H3K4me3 landscape

To explore the potential influence of S2L proteins in COMPASS-like activities, we first determined H3K4me1/2/3 global levels in *s2la* and *s2lb* lines by immunoblot analysis of chromatin extracts. No significant alterations could be detected in the *s2la-1* line. By contrast, in both Col-*0* and Nossen backgrounds *S2Lb* loss-of-function triggered a ~2-fold decrease of H3K4me3 relative to total histone H3 levels, but not of H3K4me1 and H3K4me2 enrichment (Figures 3a and Additional file 2 – Figure S5). This defect was similarly observed in *s2la-1s2lb-2* and was rescued upon complementation of the s*2lb-2* line by a *S2Lb-GFP* transgene, indicating that *S2Lb* but not *S2La* has a prominent influence on global H3-Lys4 trimethylation. Hence, we conclude that, similarly to the WDR5a AtCOMPASS core subunit [43, 44] and the SDG2 HMT [35, 36], S2Lb represents a major contributor to H3K4me3 deposition in Arabidopsis.

**Figure 3.**
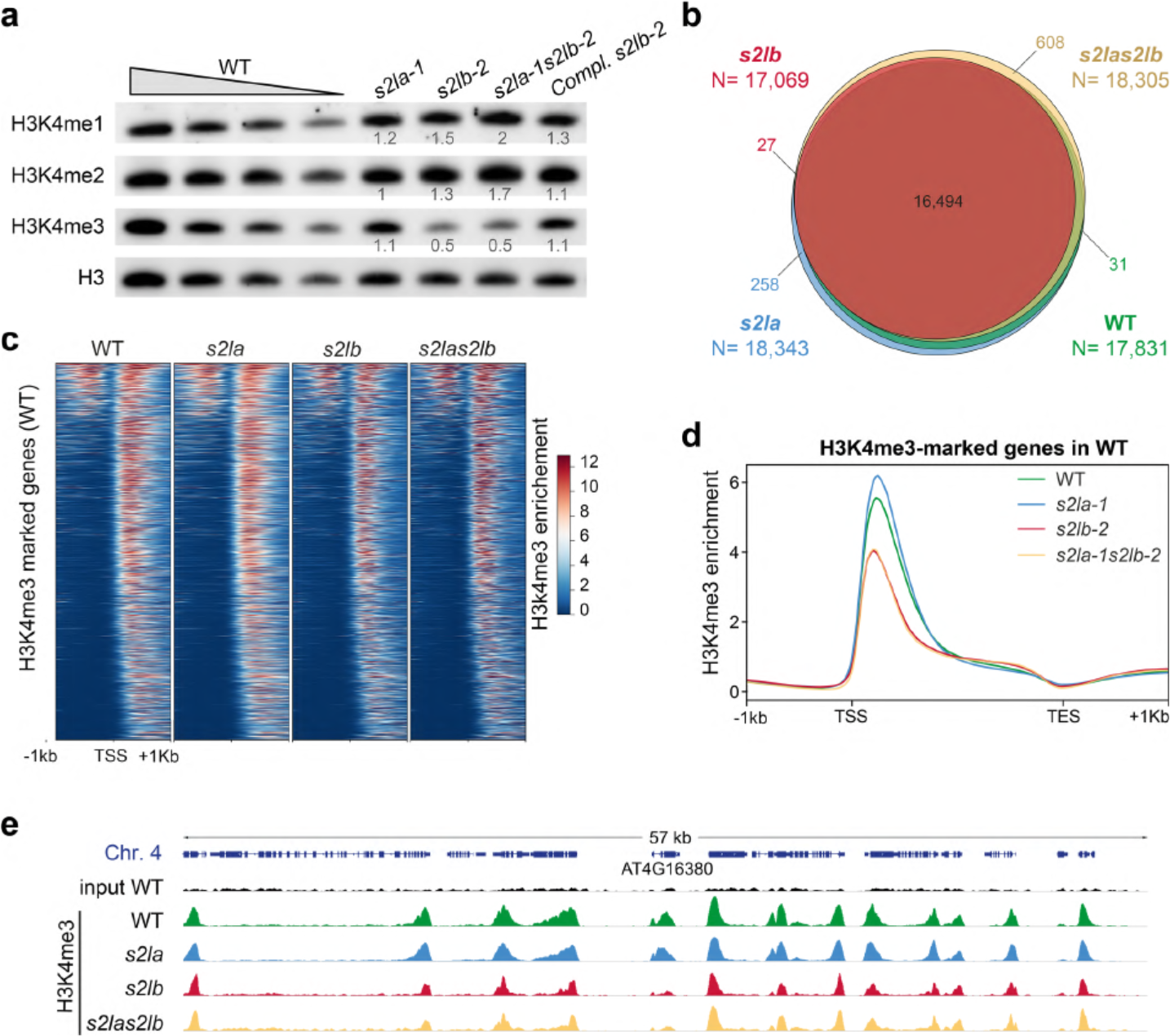
S2Lb is a major determinant of the H3K4me3 landscape. **a** Immunoblot analysis of chromatin extracts from rosette leaves of wild-type, *s2la-1*, *s2lb-2*, *s2la-1s2lb-2* and complemented *s2lb-2/S2Lb::*S2Lb-GFP plant lines. Histone H3 methylation was detected using the indicated antibodies. Decreasing quantities of chromatin extracts from wild-type plants is shown for comparison, and detection of total histone H3 is used as loading control. Quantification of H3-Lys4 methylation signals in *S2L* loss-of-function plants are given relative to their corresponding histone H3 levels. For each histone mark, levels in wild-type plants are arbitrarily set to 1. **b** ChIP-seq identification of the genes marked by H3K4me3 in WT, *s2la-1*, *s2lb-2* and *s2la-1s2lb-2* seedlings in two independent biological replicates. The corresponding gene IDs are listed in Additional file 3 (Table S2). **c** H3K4me3 enrichment over the genes marked in WT plants (N=17,831). Genes are equally ranked from top to bottom in each plant line according to the H3K4me3 median enrichment in the WT. **d** H3K4me3 median enrichment over the genes marked in WT plants (N=17,831). **e** Genome browser view of H3K4me3 enrichment over a representative region of chromosome 4. *At4g16380* is shown as an example of gene called by MACS2 as marked by H3K4me3 in WT and *s2la-1* but not in *s2lb-2* and *s2la-1s2lb-2* mutant lines. Each track represents the average data of two independent biological replicates. All tracks are equally scaled.

To determine more precisely the genomic loci impacted by S2L proteins, we first conducted a ChIP-seq analysis of the H3K4me3 landscape in six-day-old *s2la-1*, *s2lb-2* and *s2la-1s2lb-2* mutant seedlings. At this early stage, mutant and wild-type phenotypes were not visibly distinguishable. In accordance with previous H3K4me3 profiling [5, 28], H3K4me3 was mostly enriched downstream of transcription start sites (TSS) of ~18,000 genes in wild-type plants (Figure 3b-d and Additional file 2 – Figure S6).

H3K4me3 enrichment in *s2la-1* was similar to the wild-type plants in terms of the number of marked genes and the overall profile. H3K4me3 peaks could also be detected over the same repertoire of genes in *s2lb-2* and *s2la-1s2lb-2* mutants (WT-marked genes, Figure 3b), although H3K4me3 peaks are lower and/or narrower in the absence of *S2Lb* function (Figure 3d-e). Overall, these analyses showed that S2Lb, but not S2La, is required for increasing or possibly broadening H3K4me3 enrichment over most genes.

### S2Lb associates with highly transcribed genes and is required for optimal gene inducibility

To define which genes are directly targeted by S2Lb, we conducted an anti-GFP ChIP-seq analysis of a *s2lb-2*/*S2Lb::S2Lb-GFP* complemented line (4a). An EGS/formaldehyde double-crosslinking allowed us to obtain robust signals with discrete peaks (see Figure S6 and Methods) and no significant background in wild-type plants used as negative control (Additional file 2 – Figure S7). Remarkably, among the 4,557 S2Lb-GFP peaks, 97% matched H3K4me3-marked genes, altogether targeting one quarter of them (Figure 4a and Additional files 3 and 4). More precisely, profiling of reads density showed a clear tendency for co-occurrence of S2Lb-GFP and H3K4me3 in the region just downstream of TSS (Figure 4a-c). For example, S2Lb-GFP profile perfectly matched H3K4me3 domains over housekeeping genes like *TUBULIN8* but was not detected over non-expressed genes like *FT* (Figure 4c). Of note, similar S2Lb-GFP and H3K4me3 peaks of low intensity were found at different locations along the dormancy gene *DOG1*, which likely result from sense and antisense transcription start sites [57]. Also in agreement with a potential direct link between S2Lb and H3K4 trimethylation, S2Lb-GFP tend to occupy genes that are highly enriched in H3K4me3 (Figure 4d). Moreover, H3K4me3 levels over S2Lb-targeted genes were particularly decreased in the *s2lb-2* mutant line (Additional file 2 – Figure S7). Finally, we also noted that S2Lb-occupied genes typically displayed a 3′-shift of their H3K4me3 peak as compared to other H3K4me3-marked genes non-targeted by S2Lb (Figure 4d and Additional file 2 – Figure S7). These observations reveal a direct link between S2Lb and H3K4me3 enrichment over a large gene set.

**Figure 4.**
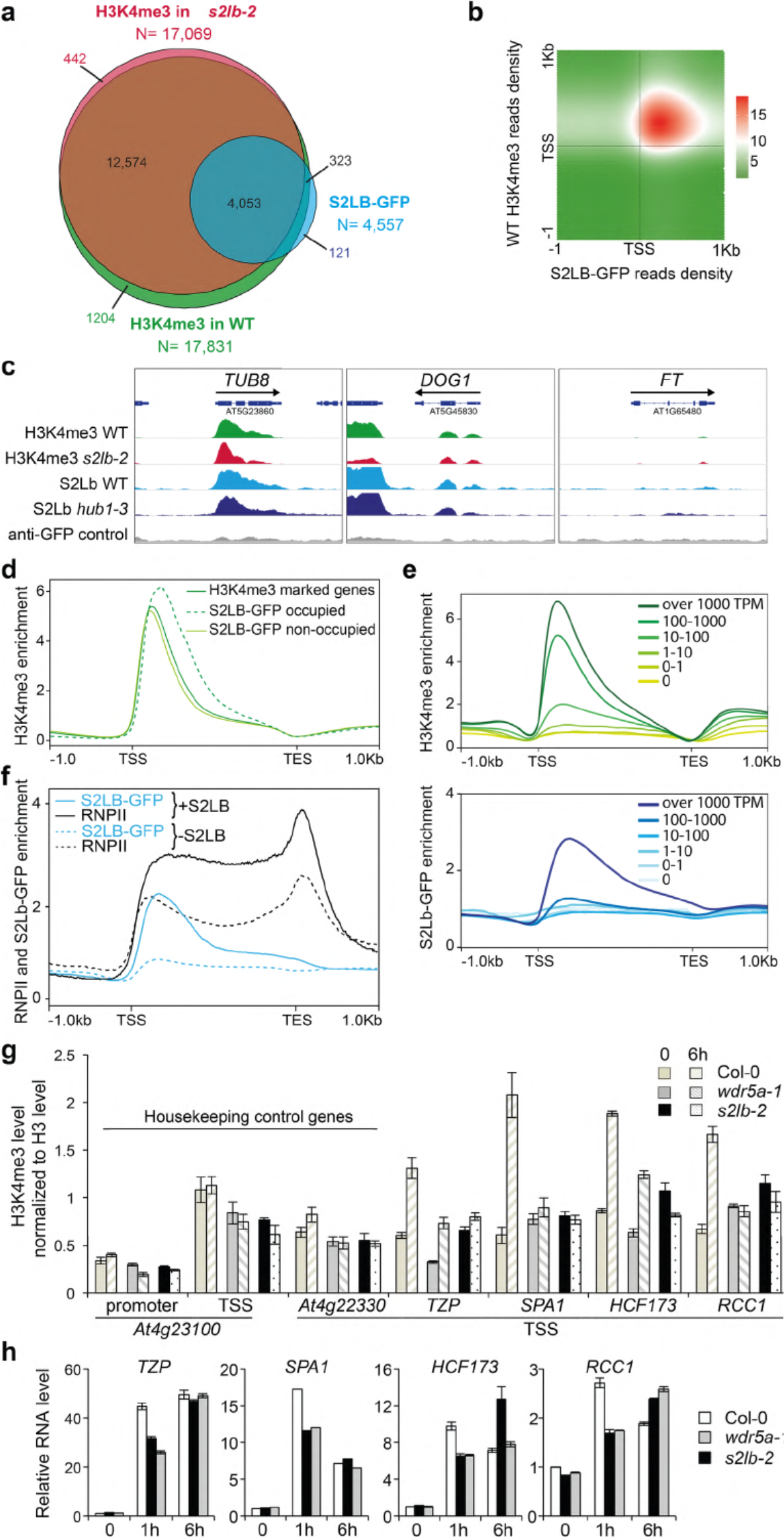
S2Lb is enriched over highly transcribed genes. **a** About 96% of the S2Lb-GFP target genes are also occupied by H3K4me3. The Venn diagram displays the overlap between genes marked by H3K4me3 in wild-type and *s2lb-2* seedlings and those occupied by S2Lb-GFP in wild-type plants. Gene IDs are listed in Additional files 3 and 4. **b** Density matrix showing the co-occurrence frequency of H3K4me3 and S2Lb-GFP along S2Lb-occupied gene annotations (N=4,557). **c** Genome browser view of H3K4me3 and S2Lb-GFP over the *TUBULIN8*, *DOG1* and *FT* genes. H3K4me3 and GFP signals in different genetic backgrounds are equally scaled. **d** Wild-type H3K4me3 median enrichment over the genes marked by H3K4me3 (dark green line; N=17,831) and those occupied by S2Lb-GFP (dashed green line; N=4,557) or not (pale green line; N=13,455). **e** Median enrichment of H3K4me3 (upper metagene plot; N=17,831) or S2Lb-GFP (lower metagene plot; N=4,557) on TAIR10 genes split into six classes ranked by expression level in wild-type seedlings. **f** Median enrichment of S2Lb and RNA Pol II over the 4,557 genes targeted by S2Lb (+S2Lb) versus the genes targeted by RNPII but not by S2Lb (-S2Lb; N=11,735). RNPII-occupied gene IDs are given in Additional file 6 - Table S5. **g** Both S2Lb and AtCOMPASS-like activities are required for H3K4me3 enrichment and induction dynamics of light-responsive genes during de-etiolation. The histogram shows H3K4me3 enrichment over several light-responsive genes in 3-day-old etiolated seedlings upon transfer from dark to light for 1 or 6h. ChIP-qPCR analyses were performed with an anti-H3K4me3 antibody and with anti-histone H3 to normalize levels to nucleosome occupancy. H3K4me3/H3 levels are given as percentages of IP/input input relative to the mean signals over two housekeeping genes (*At4g23100* and *At4g22330*) with no change in expression and in H3K4me3 level during the transition [96]. Error bars correspond to standard deviations from two replicates. **h** RT-qPCR analysis of light-responsive gene expression in samples in F). RNA levels are given relative to the wild-type dark sample (arbitrarily set to 1) normalized to *At5g13440* and *At2g36060* transcript levels. Error bars correspond to standard deviations from two replicates.

To further assess a potential link between S2Lb function and transcription, we compared wild-type and *s2lb-2* expression patterns by RNA-seq analysis. Genes misregulated in *s2lb-2* seedlings are prevalently involved in plant adaptive responses to biotic and abiotic environmental cues (Additional file 2 – Figure S8 and Additional file 5 - Table S4). In good agreement with the hypothesized influence of S2Lb on RNPII progression, *s2lb-2* misregulated genes display a biased proportion between down- and upregulated genes (60% vs 40%, respectively; Additional file 2 – Figure S8). Secondly, we assessed the relationship between S2Lb chromatin association and gene expression by quantifying its occupancy over classes of genes having different transcript levels. This showed that H3K4me3 levels correlate positively with transcript abundance while, in contrast, S2Lb-GFP targeted genes correspond to the most highly expressed genes (Figure 4e).

These observations may underscore a strict correlation between H3K4me3 marking and transcription in the bulk of different cell populations of seedlings, while in contrast S2Lb might only be detected on the most frequently transcribed genes in those various cell types. To test this hypothesis, we compared the occupancy of S2Lb-GFP and RNPII along the genome (Additional file 2 – Figure S9). RNPII ChIP-seq profiling identified about 16,000 genes that, as expected, were usually marked by H3K4me3 (88% of them) and overlapped almost entirely with S2Lb-GFP occupied genes (94% of them; Additional file 2 – Figure S9). As reported earlier in various species [8], RNPII was typically enriched along the transcribed domains, with a peak at the transcription elongation stop (TES; Figure 4e and Additional file 2 – Figure S9). Of note, RNPII enrichment was much higher on S2Lb-GFP occupied genes than on other genes (Figure 4f) as observed with H3K4me3 (Figure 4d). We concluded from these observations that S2Lb prevalently occupies genes when they are highly transcribed.

Considering these findings, the number of misregulated genes in *s2lb-2* (N=674) appears small as compared to the number of genes occupied by S2Lb-GFP (N=4,557). This contrast may suggest that H3K4me3 deposition is not sufficiently decreased upon *S2Lb* knockout to impair their transcription, or that H3K4me3 enrichment is dispensable for efficient gene expression. Another possibility is that biological deficiencies linked to H3-Lys4 trimethylation function on transcription elongation or mRNA processing would be more easily detectable during a cellular transition when genes are upregulated rather than long after reaching steady-state expression levels. To assess whether S2Lb, and more generally AtCOMPASS activity, impacts gene induction dynamics, we monitored the expression of representative light-responsive genes during de-etiolation. This morphogenic transition involves a global increase of transcription when dark-grown, etiolated, seedlings are exposed to light for the first time [58]. We tested candidate genes (*TZP*, *SPA1*, *HCF173* and *RCC1*) that were previously identified as being subject to histone H2Bub dynamics for optimal inducibility by light [59]. ChIP-qPCR and RT-qPCR showed that these genes are subject to H3K4me3 enrichment within the first 6 hours of the dark-to-light transition (Figure 4g). As expected, H3K4me3 levels displayed slower induction dynamics in *s2lb-2* seedlings and in the *wdr5a-1* RNAi line [43] than in wild-type plants. This deficiency was also true for dynamic changes in transcript levels (Figure 4h). We conclude from these analyses that, as previously proposed for H2Bub dynamics ([59], H3-Lys4 trimethylation and/or AtCOMPASS-like activity are required for optimal gene upregulation in Arabidopsis but marginally impair mRNA steady-state levels.

### S2Lb co-regulates a large set of genes with AtCOMPASS-like complexes and with SDG2

To test whether S2Lb acts as a COMPASS-like associated factor, we first assessed whether it was part of a high-molecular weight (HMW) complex by size-exclusion chromatography of soluble protein extracts from *S2Lb*::*S2Lb-GFP* plants. S2Lb-GFP eluted in two main peaks, the first one likely corresponding to its ~65 kDa monomeric form and the second one to a high-molecular weight complex of ~900 kDa or more (Figure 5a). To test whether these HMW fractions correspond to COMPASS-like complexes, S2Lb-GFP was immunoprecipitated from different fraction pools and tested for the presence of WDR5. This core subunit of AtCOMPASS-like complexes is also essential for a large fraction of H3K4me3 deposition in Arabidopsis [43–45]. S2Lb-GFP and WDR5 were both initially more abundant in monomeric fractions (Input, pool 4), but WDR5 mainly immunoprecipitated with S2Lb-GFP from HMW fractions (pools 1 and 2; Figure 5b). Hence, we conclude that S2Lb and WDR5 can associate within one or more HMW complexes *in planta*, likely corresponding to AtCOMPASS-like complexes. Association of S2Lb with WDR5a was somehow expected given the high conservation of COMPASS subunits from yeast to plants and mammals [39, 60, 61].

**Figure 5.**
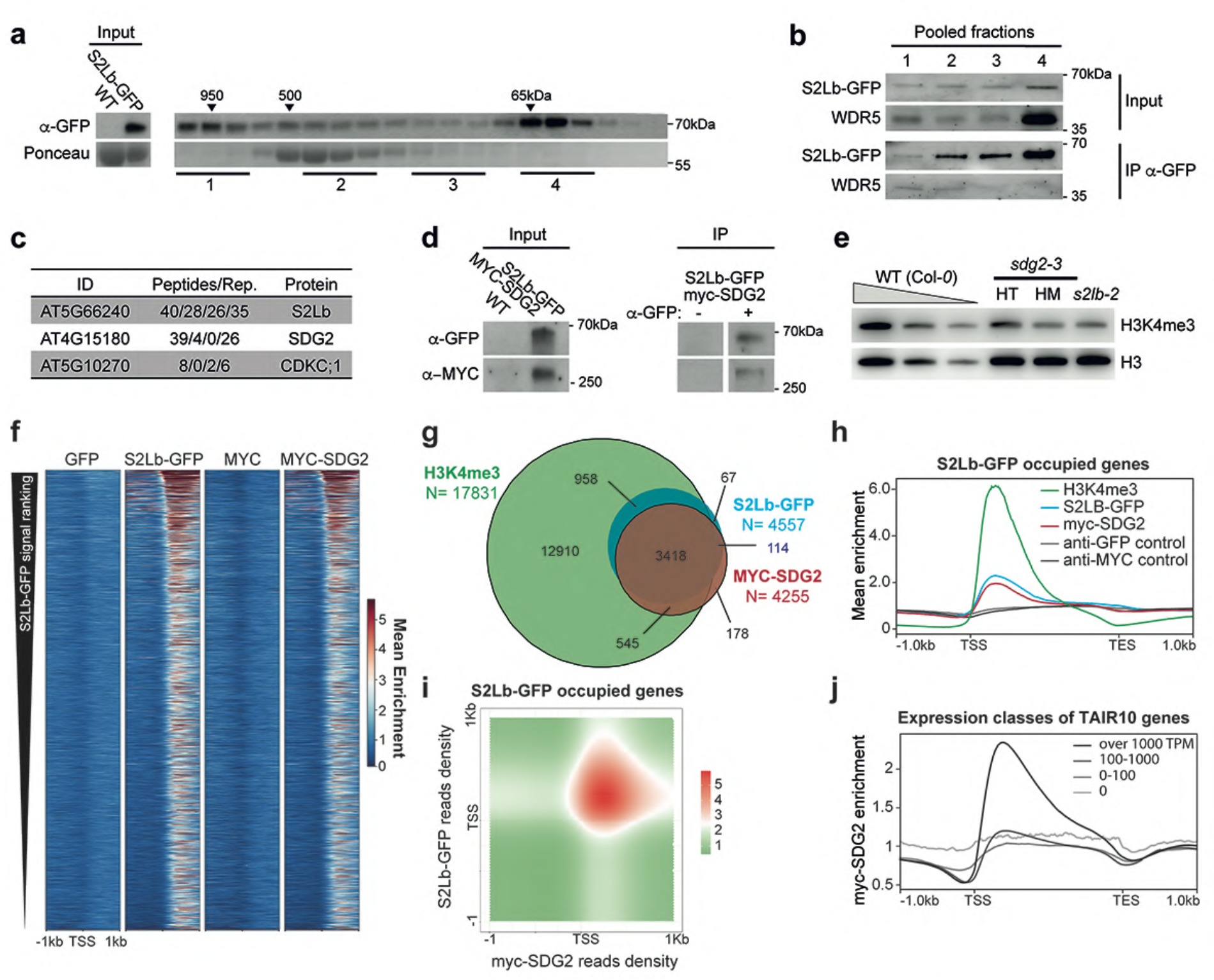
S2Lb and SDG2 associate over a large gene repertoire. **a** Size-exclusion chromatography of S2Lb-GFP complex size in *S2Lb::S2Lb-GFP* plant extracts. FPLC fractions were analyzed by anti-GFP immunoblot. Arrows indicate the elution peaks of molecular-weight standards in the same conditions. **b** Endogenous WDR5 co-immunoprecipitates with S2Lb-GFP in HMW fractions. Pooled fractions from samples in (A) were used for anti-GFP immunoprecipitation and analyzed using anti-GFP and anti-WDR5 antibodies. **c** IP-MS identification of S2Lb-GFP interacting proteins. The table gives the number of peptides from the indicated proteins detected using plant extracts from *S2Lb-GFP* plants but not from wild-type Col-*0* plants in four biological replicates. **d** Co-immunoprecipitation assay on *SDG2::MYC-SDG2;S2Lb::S2Lb-GFP* plants with anti-GFP coated (+) or uncoated beads (-). Immunoprecipitated S2Lb-GFP and MYC-SDG2 proteins were detected by anti-GFP and anti-MYC immunoblots, respectively. **e** Chromatin extracts from *S2Lb* and *SDG2* loss-of-function plants display similar defects in H3K4me3 global level. Immunoblot analysis of WT, *s2lb-2* and heterozygotes (HT) or homozygotes (HM) *sdg2-1* seedlings were performed using antibodies recognizing total histone H3 and H3K4me3. H3 is used as loading control. **f** The heatmap shows S2Lb-GFP and MYC-SDG2 enrichment over S2Lb-GFP occupied genes (N=4,557). Genes are equally ranked from top to bottom in each line according to S2LB-GFP median enrichment. **g** Overlap between genes marked by H3K4me3 and genes occupied by MYC-SDG2 or S2Lb-GFP. Overlap between S2Lb-GFP and MYC-SDG2: x5.7 pVal<0e+00; overlap between S2Lb-FP and H3K4me3: x1.7 pVal<0e+00; overlap between MYC-SDG2 ad H3K4me3: x1.6 pVal<0e+00. The corresponding gene IDs are listed in Additional files 4 and 7. **h** Median enrichment of S2Lb-GFP, MYC-SDG2 and H3K4me3 over S2Lb-GFP occupied genes (N=4,557). **i** Density matrix showing the co-occurrence of S2LB-GFP and MYC-SDG2 along S2Lb target genes (N=4,557). **j** Median enrichment of MYC-SDG2 on TAIR10 genes split into six classes ranked by expression level in wild-type seedlings.

To gain better insights into S2Lb complex activity, we conducted mass spectrometry analysis of proteins co-immunoprecipitating with GFP-S2Lb from *S2Lb*::*S2Lb-GFP* seedlings using wild-type plants as negative control. GFP-SL2b was efficiently retrieved in each of four biological replicates. Under the mild detergent conditions used, this analysis did not allow the recovery of WDR5 or other known COMPASS subunits, however the most abundantly detected protein that was robustly co-immunoprecipitated was SDG2 (Figure 5c). Of note, the CDKC;1 protein was also significantly detected in 3 out of 4 biological replicates, although with low peptide numbers. This homolog of human CDK9 belongs to the CDK9/CycT complex of P-TEFb and mediates RNPII CTD Ser-2 phosphorylation in Arabidopsis [62].

The potential association of SDG2 with S2Lb was confirmed by co-immunoprecipitation of S2Lb-GFP and MYC-SDG2 tagged proteins stably expressed in Arabidopsis under the control of their endogenous promoters (Figure 5d). Furthermore, comparison of global H3K4me3 levels in *s2lb-2* and *sdg2-3* mutant plants by immunoblot showed comparable defects (Figure 5e). Further comparison of the *s2lb-2* transcriptomic profile with all available Genevestigator datasets [52] using the Signature tool identified the *sdg2-3* profile [36] as being the most similar among all available transcriptome datasets (Additional file 2 – Figure S10). Even though distinct transcriptomic methodologies were used, direct comparison of misregulated genes in both transcriptome analyses showed that a majority of the *s2lb-2* misregulated genes display a similar trend in *sdg2-3* seedlings (55%; Additional file 2 – Figure S10).

We further examined whether S2Lb shares any functional properties with other known H3-Lys4 HMTs such as ATXR7 [33, 34] and ATX1 [32] but we identified no significant similarity with *atxr7-1* and *atx1* transcriptomic data (about 9% maximum overlap; Additional file 7 – Table S6). We conclude from these analyses that S2Lb and SDG2 directly or indirectly associate to regulate a common set of genes, possibly acting together at the chromatin level

Finally, to ascertain whether S2Lb and SDG2 co-regulate genes *in situ*, we determined the SDG2 chromatin profile by anti-MYC ChIP-seq analysis of *SDG2::MYC-SDG2* seedlings (Figure 5f and Additional file 9 - Table S8). This unveiled that SDG2 associates with a similar number of chromatin loci than SL2b (4,255 vs 4,557), 80% of them occurring on the same genes (Figure 5g). Metagene-plot analyses showed very similar enrichment profiles over gene bodies (Figure 5h), mainly co-occurring over 3′ domains of TSS (Figure 5i). As described above for S2Lb-GFP, almost all genes occupied with MYC-SDG2 were marked by H3K4me3 (Figure 5g) and tend to be highly expressed (Figure 5j). Searching for potential DNA-sequence contexts underlying S2Lb and/or SDG2 association with chromatin, we were not able to identify a specific set of transcription factor binding motifs, apart from diverse forms of GA stretches (Additional file 8 – Table S7). Collectively, these findings show that S2Lb associates with one or more AtCOMPASS-like complexes and co-regulate multiple genes in association with the SDG2 histone methyltransferase.

### S2Lb chromatin association and H3K4me3 deposition do not depend on prior histone H2B monoubiquitination

The yeast Swd2 homolog of S2Lb is thought to drive yCOMPASS-mediated H3K4me3 deposition on H2Bub-modified nucleosomes [14–16]. The apparent conservation of SWD2-Like proteins in Arabidopsis may suggest that S2Lb is also required for H3K4me3 deposition or maintenance through a similar mechanism of action. Accordingly, genome profiling further showed a clear tendency of S2Lb-GFP to occupy H2Bub-marked genes (Figure 6a), suggesting that some H2Bub-enriched domains allow recruitment of AtCOMPASS activity. Nevertheless, differently from yeast, the bulk of H3K4me3 is maintained in mutant plants lacking or over-accumulating H2Bub (Figure 6b and [63–67]). Vice versa, H2Bub levels are not visibly affected in *s2lb-2* and *sdg2-3* plants defective in H3K4me3 deposition (Figure 6c).

**Figure 6.**
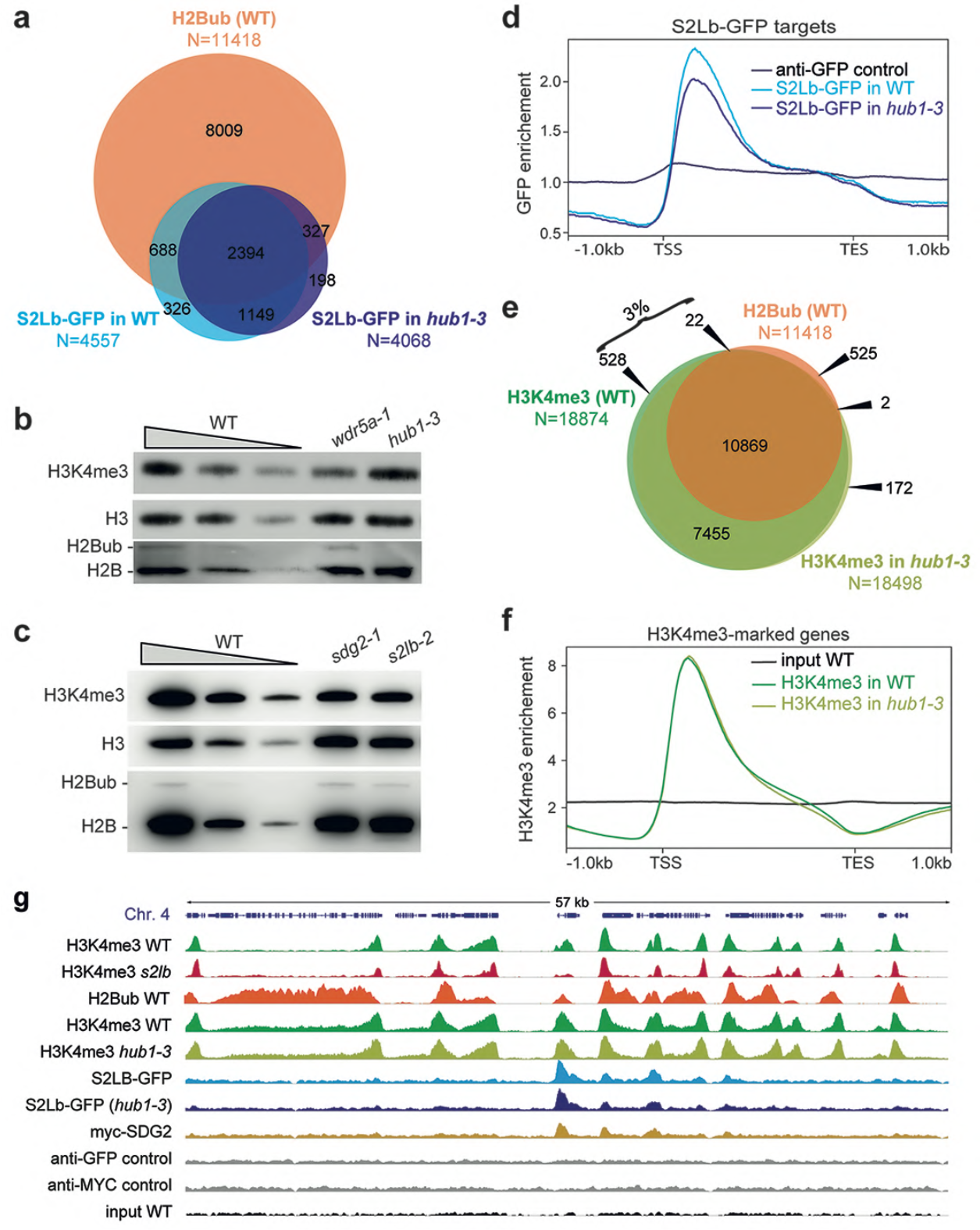
S2Lb chromatin association and H3K4me3 deposition are usually independent from histone H2B monoubiquitination. **a** Overlap between genes marked by H2Bub (from [95]) and by S2LB-GFP in wild-type (WT) or in *hub1-3* seedlings. For proper comparison of S2Lb-GFP enrichment, WT/*S2Lb::S2Lb-GFP* and *hub1-3*/*S2Lb::S2Lb-GFP* seedlings were grown and processed for ChIP-seq in parallel. The corresponding gene IDs are listed in Additional file 4 (Table S3). **b** Wild-type H3K4me3 levels in chromatin extracts of *HUB* loss-of-function plants. Decreasing quantities of chromatin extracts from wild-type plants is shown for comparison, and detection of total histone H3 and H2B forms are used as loading controls. **c** Wild-type H2Bub levels in *S2Lb* and *SDG2* loss-of-function plants. Immunoblots were performed as in b. **d** S2Lb-GFP median enrichment in WT/*S2Lb::S2Lb-GFP* and *hub1-3*/*S2Lb::S2Lb-GFP* seedlings on the S2Lb-GFP targeted genes (N=4,557). **e** Overlap between H2Bub-marked genes (from [95]) and H3K4me3-marked genes in wild-type (WT) or in *hub1-3* seedlings (this study). For proper comparison of H3K4me3 marking, wild-type and *hub1-3* seedlings were grown and processed for ChIP-seq in parallel, independently from other ChIP-seq in Figures 3 to 5. The corresponding gene IDs are listed in Additional file 9. **f** H3K4me3 median enrichment in WT and *hub1-3* seedlings on the genes marked by H3K4me3 in WT (N=18,874). **g** Genome browser view of H3K4me3, S2Lb-GFP and MYC-SDG2 over the same region of chromosome 4 shown in Figure 3. H3K4me3 and GFP or MYC signals in different genetic backgrounds are equally scaled.

These observations suggest an absence or at most only limited AtCOMPASS-mediated H2Bub/H3K4me3 crosstalk in Arabidopsis, and further interrogates 1) whether H3K4me3 patterns along the genome rely, even partially, on histone H2B monoubiquitination and 2) how S2Lb, AtCOMPASS and SDG2 are recruited to chromatin loci.

To address these questions, we took advantage of *hub1-3* mutant plants in which the deposition of histone H2B monoubiquitination is abolished to test whether S2Lb-GFP recruitment and H3K4me3 enrichment rely on H2Bub. H2Bub levels are undetectable in homozygous *hub1-3* seedlings ([48, 49, 59] and Figure 6b). Upon introgression of *S2Lb::S2Lb-GFP* in the *hub1-3* background, ChIP-seq analysis of S2Lb-GFP showed that about one third of the SL2b-targeted genes were different in the *hub1-3* plants, with a tendency to be marked in WT but not in *hub1-3* background (1,014 vs 525 genes; Figure 6a). SL2b-GFP enrichment over gene bodies was slightly decreased in *hub1-3* plants, still with a similar profile (Figure 6d). This analysis showed that S2Lb-GFP can be recruited over many H2Bub-marked genes largely independently of this histone, whereas a minority of genes might be subjected to a COMPASS-based histone crosstalk. To test this second hypothesis, we compared the set of genes that lost both S2Lb-GFP occupancy and H3K4me3 marking in *hub1-3* mutant plants. Only nine genes corresponding to this criterion could be identified (Additional file 2 – Figure S11). We concluded that S2Lb-GFP and H3K4me3 may aberrantly target other genes in *hub1-3* plants, possibly as a consequence of mild transcriptomic and phenotypic variations induced by *HUB1* loss-of-function [48, 49, 59].

Direct comparison of H3K4me3 profiles in wild-type and *hub1-3* seedlings further showed that only ~3% of the genes lose H3K4me3 enrichment in the absence of H2Bub (Figure 6e and Additional file 10 - Table S9). Moreover, contrasting with PAF1c mutant plants [68], global H3K4me3 enrichment and positioning along the 5′ domains of gene bodies were not detectably affected by loss of HUB activity (Figure 6f and Additional file 2 – Figure S12), again supporting that the vast majority of genes are subject to H3-Lys4 trimethylation independently from H2Bub deposition.

The genome-wide profiles obtained in this study confirmed the spatial correlation between S2Lb-GFP and MYC-SDG2 peaks over about one third of the H3K4me3-marked genes (Figure 6g). H3K4me3 enrichment was robustly diminished in the *s2lb-2* line while, in contrast both H3K4me3 and S2Lb-GFP occupancy were typically unaffected in the *hub1-3* line (Figure S6). Collectively, we conclude that AtCOMPASS and SDG2 mainly drive H3-Lys4 trimethylation through H2Bub-independent pathways in Arabidopsis.

## Discussion

### S2Lb and COMPASS-like proteins as partners of the plant-specific histone methyltransferase SDG2

In this study, we first report that S2Lb, a homolog of the yCOMPASS-associated subunit, is a major actor of H3-Lys4 trimethylation with SDG2 despite the absence of clear H2Bub-H3K4me3 histone crosstalk in Arabidopsis. Accordingly, GFP-tagged S2Lb resides only on H3K4me3-enriched genes, mostly those displaying the H3K4me3 HMT protein SDG2 [35, 36]. ChIP-seq analyses further showed a tight co-occurrence of S2Lb-GFP with MYC-SDG2 over thousands of genes, in particular those displaying high RNPII occupancy and elevated transcript levels. Our ChIP analyses do not allow us to ascertain whether S2Lb, SDG2 and RNPII physically associate onto the same chromatin fragments, but in support of this possibility SDG2 was the most abundantly co-purifying protein with S2Lb.

A series of complementary evidence points towards a functional partnership between S2Lb and SDG2 with one or more COMPASS-like complexes in Arabidopsis. Firstly, S2Lb and SDG2 display very similar genomic distributions, by co-occuring just downstream from the TSS of more than four thousand H3K4me3-marked genes. The reciprocal comparison is not true as the majority of H3K4me3 peaks do not overlap with S2Lb or SDG2 peaks. S2Lb and SDG2 might only be detected over the genes most frequently transcribed among the various seedling cell types or formerly transcribed genes might retain H3K4me3 marking but not S2Lb association. Secondly, S2Lb and SDG2 are both important for establishing or maintaining H3K4 trimethylation levels since their loss-of-function leads to a similar 50-70% decrease of the bulk of H3K4me3 [35, 36]. Thirdly, *S2Lb* and *SDG2* loss of function plants share several related phenotypic defects throughout the life cycle including dwarfism, short roots, loss of apical dominance and impaired fertility [35, 36, 69, 70]. Accordingly, transcriptomic analyses revealed a striking overlap between genes misregulated in *s2lb-2* and in *sdg2-3*, indicating that S2Lb and SDG2 regulate expression of a common gene set.

IP-MS attempts to recover WDR5 or other known COMPASS subunits by IP/MS of GFP-S2Lb were unsuccessful, possibly because of the mild detergent conditions used in these assays as a consequence of the large MYC-SDG2 protein (~300 kDa-migrating form) being largely insoluble in plant extracts. Notwithstanding, S2Lb and WDR5 successfully pulled-down from one or more high-molecular weight complexes from soluble plant extracts. They presumably interact indirectly as shown for yeast Swd2 and Swd3 [71]. Their association was somehow expected given the high conservation of COMPASS subunits from yeast to plants and mammals [39, 60, 61], but less so for a plant-specific protein like SDG2. We have been able to identify a significant relationship between the misregulated gene repertoire in *s2lb* seedlings and *SDG2* loss-of-function seedlings but less for other H3K4me3 HMTs such as *ATX1* and *ATXR7*. S2Lb therefore seems to have certain level of specificity for SDG2, which may relate to the wide expression pattern of these two genes throughout plant development. Hence, the evolutionarily conserved COMPASS-like complexes not only act with Trithorax-like proteins such as ATX1, SDG14 and SDG16 in Arabidopsis as in other species [43, 44] but has also been coopted by the plant-specific SDG2 HMT (Figure 7).

**Figure 7.**
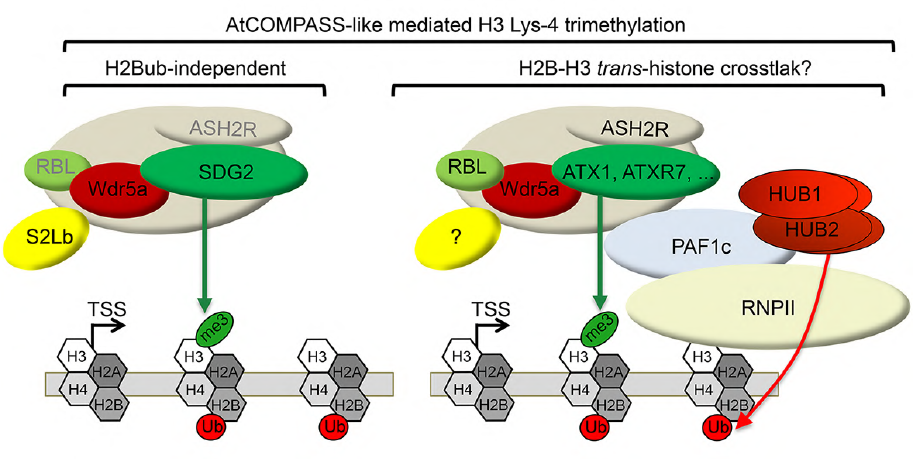
Working model for S2Lb activity with AtCOMPASS-like and SDG2 in histone H3-Lys4 trimethylation. In a major pathway, S2Lb and SDG2 target highly transcribed genes for H3-Lys4 trimethylation as part of a COMPASS-like complex acting independently of H2B monoubiquitination. S2Lb might as well be part of other COMPASS-like complexes with other histone methyltransferases such as ATX1 and ATXR7. Arabidopsis WDR5a, RbBP5-Like (RBL) and Ash2-related (ASHR) are core components of AtCOMPASS-like complexes [37, 43–45] but the nature of their physical association with SDG2 remains to be determined. SDG2 might additionally act independently from S2Lb in different genic or cellular contexts, possibly with S2La. Whether histone H2B monoubiquitination is required for other types of At-COMPASS complexes activity and acts co-transcriptionally with PAF1c as in other species also remains to be resolved.

Observation that both *SDG2* and *WDR5* null mutations are sterile whilst mutant plants combining *S2la* and *S2Lb* knockout are partially fertile suggests that S2L proteins are less essential than SDG2 and AtCOMPASS complexes. SDG2 may either catalyze H3K4 trimethylation alone or with COMPASS-like independently from S2Lb, potentially having a residual activity at specific chromatin loci or in specific cell types such as in male and/or female gametophytes. Whether S2Lb physically interacts with SDG2 remains to be resolved, as does the question of whether SDG2 associates with WDR5a within an AtCOMPASS super-complex with other histone modifying and remodeling activities, as recently identified for the AtCOMPASS-FRIGIDA complex [72].

### H2Bub-independent AtCOMPASS-like activity

Several independent studies have revealed that the bulk of H3K4me3 is retained in mutant plants lacking or over-accumulating H2Bub and in Paf1c mutants [68], as reproducibly shown by immunoblot analyses here and by several others ([63–67] and Figure 6b). Targeted ChIP-qPCR has also been conducted over a handful of genes in *hub* mutant plants, such as the flowering regulatory genes *FLC* [63], *SOC1*, *FT* and *MAF4* [67] and the clock component genes *CCA1*, *TOC1* and *ELF4* [73], showing in all cases that H3K4me3 level was lower than in wild-type plants. Genetic approaches combining mutations impairing H2B monoubiquitination and histone methylation identified both additive and synergistic effects on Arabidopsis phenotypic quantitative traits, suggesting the existence of interplays among different histone modifications [67]. Still, the lack of mechanistic assessments and of genome-wide resolution have not allowed an unambiguous evaluation of whether an H2Bub-H3K4me3 *trans*-histone crosstalk is at play in plants. Here, we first observed using two independent seed batches and upon certifying that homozygous seeds were used, that H3K4me3 profiles were quasi indistinguishable between wild-type plants and *hub1-3* mutant plants lacking detectable H2Bub. Using other harvesting daytime, growth conditions or developmental stages might possibly be more accurate to comparing our results with former studies. Nevertheless, considering that Swd2 allows tethering COMPASS on H2Bub-modified nucleosomes in other species [14–16], a second approach consisted in assessing whether S2Lb is recruited onto the epigenome in the absence of H2B monoubiquitination as a proxy to test for a potential AtCOMPASS-mediated H3K4me3-H2Bub crosstalk. Although enrichment levels were slightly weaker, the vast majority of S2Lb target genes were occupied by S2Lb-GFP in both wild-type and *hub1-3* seedlings. Hence, our two complementary approaches point towards a role for AtCOMPASS/SDG2 in driving H3-Lys4 trimethylation that is largely independent from histone H2B monoubiquitination in Arabidopsis.

A 3′-shift of the H3K4me3 peak was observed on S2Lb-targeted genes. A similar defect has been reported in Arabidopsis PAF1c mutant seedlings, and proposed to result from an irregular transition from the Ser-5 to Ser-2 phosphorylated form of RNPII [68]. Considering that most genes were still marked by H3K4me3 in *S2Lb* mutant plants, albeit to a lower extent, we propose that S2Lb is required for maintenance or broadening of the H3K4me3 landscape during RNPII transition into productive elongation while it might not be involved in its nucleation. If true, this hypothesis would provide a rationale for conserving SWD2-like activities in plants despite not contributing to a recognizable *trans*-histone crosstalk function.

These findings add to our former report that H3K4me3 is efficiently established over light-responsive genes in *hub1-3* seedlings during upon their induction [59]. In the absence of H2Bub-H3K4me3 *trans*-histone crosstalk, AtCOMPASS complexes might rather be recruited onto chromatin loci in a sequence-specific manner and in response to specific signals by means of transcription factors. A targeting mechanism by transcription factors such as bZIP28 and bZIP60 has recently been unveiled for the regulation of endoplasmic reticulum stress responsive genes [47].

### Complex relationships between histone H3 Lys-K4 trimethylation and histone H2B monoubiquitination with transcription regulation in Arabidopsis

RNA-seq analysis showed that only a small subset of H3K4me3-marked and of S2Lb-targeted genes were misregulated in young *S2Lb* knockout seedlings. This is line with the apparent wild-type phenotype of *s2lb* seedlings at this early developmental stage but also appears counter intuitive with the proposed role of S2Lb in AtCOMPASS activity and with the instructive role of H3 Lys-4 trimethylation on RNPII processivity. Still, as for *S2Lb*, both *HUB* and *PAF1c* loss-of-function trigger weak transcriptomic defects in Arabidopsis [49, 63, 68]. Depletion of H3K4me3 has only marginal effects on gene expression in other species as well [12]. Hence, H3K4me3 may contribute to the reinforcement of the active state of transcription [68] and to the fine-tuning of genome expression during plant development and adaptive responses [67], a proposed function that is consistent with our observation that AtCOMPASS-like deficient plants are impaired in the accurate inducibility of light-regulated genes. Investigating more precisely the effect of S2Lb or other COMPASS subunits on transcription efficiency would probably require a quantification of nascent transcripts production in a dynamic system such as de-etiolation or another cellular adaptive response.

The CDKC;1 protein was detected as co-purifying with S2Lb in our IP-MS analyses, although not systematically and with low peptide numbers. This association might be functionally meaningful, as CDKC;1 mediates RNPII CTD Ser-2 phosphorylation in Arabidopsis [62, 74–76] and acts as an activator of transcription in plants [77]. CDKC;2, another cyclin-dependent kinase involved in RNPII regulation has also been found recently to co-purify with HMTs and chromatin remodeling factors using similar approaches [78]. Such interactions could potentially link S2Lb directly to the regulation of RNPII CTD phosphorylation and therefore to the transition towards transcription elongation.

### A diversity of COMPASS-like complexes in *A. thaliana*

*A. thaliana* encodes two paralogs of the *S. cerevisiae SWD2* gene with similar expression patterns in most organs. *S2La* is usually expressed to a much lower level than *S2Lb*, possibly targeting only a few genes or acting in a few cell types. This is also the case for *ATX1* [35, 79], which presumably targets a few specialized genes on which it helps recruiting a COMPASS-like complex and promotes assembly of the RNPII pre-initiation complex [37]. Both *S2L* genes encode euchromatic proteins that differ in their structural structure. *S2La* disruption does not detectably aggravate either *s2lb-2* morphological phenotypes or its H3K4me3 defects. Hence, S2La could also allow for a distinct, minor, H3K4me3 deposition pathway, possibly acting in a *trans*-histone crosstalk with H2B monoubiquitination on specific genes, or rather act in other histone modifications.

S2La and S2Lb polymorphic WD40 repeat domains may underpin different protein association capacities, for example influencing their association with different transcription factors targeting SDG2 or other HMTs to distinct loci, or with other protein complexes. Noteworthy, yeast SWD2 is also an integral subunit of the Cleavage and Polyadenylation Factor (CPF) complex involved in 3′end mRNA processing [80, 81]. In agreement with its role in H3K4me3 deposition, predominant phenotypes induced by *S2Lb* loss-of-function are shared with COMPASS [40, 43–45] and H3K4 trimethylation defective phenotypes (SDG2), most notably dwarfism, impaired fertility and early flowering [35, 36]. In contrast, *s2la-1* plants are late flowering like CPF subunit mutants [82]. Hence, two *SWD2* paralogs might be specialized in Arabidopsis, a situation previously identified in *S. pombe* [83]. Given the ancient origin of the duplication event of *S2La* and *S2Lb* in the plant lineage, examination of their functional diversification represents an interesting aspect to decipher in future studies.

### Conclusion

By contrast with *S. cerevisiae* in which a single SET1 protein catalyzes histone H3 Lys-4 trimethylation as part of COMPASS acting upon histone H2B monoubiquitination, in Arabidopsis H3K4me3 deposition is mediated by multiple ubiquitous or cell-specific histone methyltransferases (HMT). Here, we show that a major pathway for H3 Lys-4 trimethylation involves the plant specific HMT SDG2 acting in the context of an evolutionarily conserved COMPASS-like activity in Arabidopsis. In addition, we report that a Swd2-like (S2Lb) COMPASS axillary subunit is recruited onto most transcribed genes along with SDG2 and allows increasing H3K4me3 occupancy in wild-type plants but also in plants lacking H2Bub. Collectively, this study sheds light on the evolution of SWD2-like proteins and COMPASS-like activity, which might underpin an atypical and H2Bub-independent pathway driving most H3K4me3 deposition in plants.

## Methods

### Plant material and growth conditions

All Arabidopsis lines used in this study are in the Col-*0* background except *s2lb-1* and its parental line Ds1-388-5 that are in a Nossen background. The *s2la-1* T-DNA insertion line (WiscDsLox489-492K11) described in [50] was obtained from NASC [84]. The *s2lb-2* line (RATM54-3645-1) was obtained from the RIKEN Institute [85] and subjected to five successive backcrossing with Col-*0* wild-type plants as female counterparts to generate *s2lb-2* plants. The *wdr5a-1* RNAi line and the *sdg2*/SDG2::myc-SDG2 line have previously been described [36, 43]. Plants were grown under 100 µmol.m².s^−1^ light in soil or *in vitro* under long-day (16h day 23°C/8h night 19°C) conditions (except for the indicated flowering time experiments). For *in vitro* growth, seeds were surface sterilized and plated on MS medium containing 0.9% agar and stratified for 3 days at 4°C before transfer to growth chambers. Root length was determined on seedlings grown *in vitro* on vertical MS plates supplemented with 1% sucrose. Position of root tips was marked every two days from day 3 to day 11 post-germination. Plates were scanned at day 11 and root length was measured using Image J [86]. Deetiolation experiments were conducted as in [59].

### Dormancy assay

Dormancy was measured on seeds issued from 3 independent productions after plant growth at 20-22°C under a long-day photoperiod. At full maturity, seeds were harvested and germination was assessed at 15°C and 25°C in darkness in 3 biological replicates of 50 seeds for each genotype. Experiments were conducted in 9 cm Petri dishes on a layer of cotton wool covered by a filter paper sheet soaked with water. A seed was considered as germinated when the radicle has protruded through the testa. Germination was scored daily for 10 days and the results presented correspond to the mean of the germination.

### Plasmid construction

The *p35S*::GFP-S2La construct was generated by inserting the entire coding sequence of *S2La* (including stop codon) amplified from wild-type Col-*0* cDNA downstream of the GFP coding sequence in the pB7WGF2 plasmid (Ghent plasmids collection, http://bccm.belspo.be/index.php) via Gateway technology (Invitrogen). The same was done for the *p35S*::GFP-S2Lb construct, except that the entire coding sequence of *S2Lb* was obtained from the U16729 pENTR-D-TOPO plasmid (ABRC). The

*pS2Lb*::S2Lb-GFP and *pS2Lb*::S2Lb constructs were generated by inserting a PCR-amplified 3.1 kb *S2Lb* genomic fragment (entire genomic coding sequence and 1 kb of promoter region) in frame with a downstream *GFP* reporter gene in the pB7FWG,0 plasmid, or in the pB7WG plasmid respectively (Ghent plasmids collection, http://bccm.belspo.be/index.php) via Gateway technology (Invitrogen). As the *S2Lb* fragment was cloned without STOP codon, a TAG codon was then introduced in the *pS2Lb*::S2Lb construct by changing one nucleotide using a site-specific mutagenesis kit (QuikChange XL Site-directed mutagenesis kit, Agilent).

### *In situ* immunolocalization

Five-day-old wild-type, *p35S*::GFP-S2La, *p35S*::GFP-S2Lb and *pS2Lb*::S2Lb-GFP seedlings were vacuum infiltrated in 4% formaldehyde, 10 mM Tris-HCL pH 7.5, 10 mM EDTA, 100 mM NaCl for 30 min and washed with Tris buffer. Cotyledons were chopped in ice-cold LB01 buffer (15 mM Tris-HCl at pH 7.5, 2 mM EDTA, 0.5 mM spermine, 80 mM KCl, 20 mM NaCl, 0.1% Triton X-100) and nuclei were isolated using a Douncer (Wheaton), filtered through a 50-µm nylon mesh, centrifuged at 500g for 5 min at 4°C, spread and air dried on APTES/glutaraldehyde treated slides. Slides were post-fixed in methanol-acetone 1:1 solution for 10 min and blocked in PEMSB for 2h at room temperature. The slides were incubated overnight at 4°C with a primary antibody specific to GFP (1/200, Life Technologies, A-11122) then for 2 hours with goat-anti-rabbit-AlexaFluor488 secondary antibody (Life Technologies, A-11008). Slides were washed and mounted in Vectashield with 2µg/µl DAPI. Images were taken using a confocal laser-scanning microscope (SP5, Leica).

### Immunoblot analyses

Soluble protein samples were obtained using the indicated methods, and chromatin extracts were obtained as previously described [59]. Unless stated, ten micrograms of protein samples were loaded on 14% LiDS Tris-Tricine gels and blotted onto PVDF membranes before immunodetection and analysis using a LAS4000 luminescence imager (Fuji). The following antibodies were used: anti-H3 (Millipore #07-690), anti-H3K4me1 (Active Motifs #39297), anti-H3K4me2 (Millipore #07-030), anti-H3K4me3 (Millipore #05-745), anti-GFP (Clontech #632381), anti-MYC antibodies (Millipore #05-724) or custom-designed anti-rice histone H2B [59]. Anti-WDR5 serum was obtained by immunization of a rabbit with a 50-amino-acid synthetic peptide corresponding to amino acids 42-91 of the WDR5a protein and affinity purification by the SDIX company (USA).

### Gel filtration

Size exclusion chromatography was performed as previously described [87]. Elution fractions were either analyzed by immunodetection of S2Lb-GFP on 40µl samples or pooled as indicated before immunoprecipitation using a Pierce Crosslink IP kit (Thermo Scientific) and an anti-GFP antibody (Molecular Probes #A-11122).

### *In vivo* pull-down assays

*In vivo* pull-down assays were performed on 1 mg of protein extracts from 10-day-old seedlings. Proteins from *pSDG2::myc-SDG2*/*pS2Lb::S2Lb-GFP* homozygous plants obtained by crossing the two respective lines were extracted using a modified RIPA buffer (Tris pH 7.6 25 mM, NaCl 150 mM, NP40 1%, Sodium deoxycholate 1%, SDS 0.1% and protease inhibitors). After clearing the samples with uncoupled beads (ChIP Adembeads, Ademtech), S2Lb-GFP proteins were immunoprecipitated for 2 hours using a GFP-trap system (Chromotek) coupled to magnetic beads. A mock was done using uncoupled beads. Beads were washed with RIPA buffer without detergents before elution with 2X Laemmli buffer. Eluates were analyzed on 8% SDS-PAGE gels and blotted onto PVDF membranes before immunodetection.

### SDG2 and S2Lb affinity purification and mass spectrometry

For each biological replicate, protein samples were immuno-isolated from 2g of either wild-type, *pSDG2::myc-SDG2*, or *pS2Lb::S2Lb-GFP* 10-day-old seedlings as described above using either GFP-or MYC-TRAP slurries (Chromotek # gtma-20 and #yta-20, respectively) in modified RIPA buffer to allow for SDG2 affinity purification. For mass spectrometry, SDS/PAGE was used without separation as a clean-up step, and only one gel slice was excised. Gel slices were washed and proteins were reduced with 10 mM DTT before alkylation with 55 mM iodoacetamide. After washing and shrinking the gel pieces with 100% (vol/vol) MeCN, in-gel digestion was performed using trypsin/LysC (Promega) overnight in 25 mM NH_4_HCO_3_ at 30 °C. Peptides were analyzed by LC-MS/MS using an RSLCnano system (Ultimate 3000, Thermo Scientific) coupled to an Orbitrap Fusion Tribrid mass spectrometer (Thermo Scientific). Peptides were loaded onto a C18-reversed phase column (75-μm inner diameter × 2 cm; nanoViper Acclaim PepMap^TM^ 100, Thermo Scientific), separated and MS data acquired using Xcalibur software. Peptides separation was performed over a linear gradient of 100 min from 5% to 30% (vol/vol) acetonitrile (75-μm inner diameter × 50 cm; nanoViper C18, 2 μm, 100Å, Acclaim PepMap^TM^ RSLC, Thermo Scientific). Full-scan MS was acquired in the Orbitrap analyzer with a resolution set to 120,000 and ions from each full scan were HCD fragmented and analyzed in the linear ion trap. For identification the data were searched against the *Arabidopsis thaliana* TAIR10 database (2016) using Mascot 2.5.1 (Matrix Science). Enzyme specificity was set to trypsin and a maximum of two-missed cleavage sites were allowed. Oxidized methionine, N-terminal acetylation, and carbamidomethyl cysteine were set as variable modifications. Maximum allowed mass deviation was set to 10 ppm for monoisotopic precursor ions and 0.4 Da for MS/MS peaks. The resulting files were further processed using *my*ProMS v3.0 [88]. FDR calculation used Percolator and was set to 1% at the peptide level for the whole study. Unless indicated otherwise, a protein was considered present if at least three peptides in all three biological replicates were detected for qualitative analysis of immuno-isolated samples.

### Protein sequence analyses

Full-length protein sequences were aligned with ClustalW using default parameters. The alignment was used to construct a neighbor-joining tree using MEGA4. Bootstrap values were obtained after 1,000 permutation replicates. WD40 repeats were determined using the WDSP predicting software [89, 90].

### RT-PCR analyses

For seedlings, total RNA was extracted using NucleoSpin RNA Plant (Macherey-Nagel). For seeds, 70 mg aliquots of seeds were ground in liquid nitrogen, and total RNA was extracted using a modified CTAB method [91]. Reverse transcription and subsequent quantitative PCR were performed on 1 µg of DNaseI-treated (Invitrogen, Amplification Grade DNaseI) RNAs using random hexamers and a cDNA reverse transcription kit (Applied Biosystems). Quantitative PCR was performed using LightCycler 480 SYBR green I Master mix and a LightCycler 480 (Roche). To confirm absence of contamination of the samples by genomic DNA, PCR was also performed using primers flanking one intron of *ACTIN2* and the size of the amplicons was checked on agarose gels. Data were normalized relative to genes with invariable expression as indicated in figure legends.

### RNA-sequencing and bioinformatics

Wild-type Col-*0* and *s2lb-2* seedlings were grown *in vitro* under long-day conditions and harvested 6 days after germination at 8 ZT. Two independent biological replicates for each genotype were produced using different seed batches. Total RNA was extracted using NucleoSpin RNA Plant (Macherey-Nagel). Messenger (polyA+) RNAs were purified from 1 µg of total RNA using oligo(dT). Libraries were prepared using the strand specific RNA-Seq library preparation TruSeq Stranded mRNA kit (Illumina). Libraries were multiplexed by 4 on 1 flowcell lane. A 50-bp single read end sequencing was performed on a HiSeq 1500 device (Illumina). A minimum of 37 million passing Illumina quality filter reads was obtained for each of the 4 samples. TruSeq adapters were removed with trimmomatic v0.36 [92] using the parameters “ILLUMINACLIP:TruSeq3-SE.fa:2:30:10 LEADING:5

TRAILING:5 MINLEN:20”. Reads were mapped on TAIR10 genome assembly of *A. thaliana* genome providing the gene annotation obtained from Araport11 [93] using STAR [94]. The command used is “STAR --genomeDir STAR_2.5.4b.TAIR10 -- quantMode GeneCounts --outSAMstrandField intronMotif --sjdbOverhang 100 --sjdbGTFfile Araport11_GFF3_genes_transposons.201606.gtf -- outSAMtype BAM SortedByCoordinate -- outFilterIntronMotifs RemoveNoncanonical”. Differentially expressed genes were identified with DESeq2 (adj.p-value < 0.01). Genes were split in 4 groups based on the normalized read counts in wild-type (TPM equal to 0, TPM from 0 to 100, TPM from 100 to 1000, TPM above 1000) and used for Figure 5j.

### ChIP-qPCR, ChIP-sequencing and ChIP bioinformatics

Plants were grown *in vitro* under long-day conditions and whole seedlings were harvested 6 days after germination at 8 ZT. Chromatin extraction and immunoprecipitation of histones were performed as previously described [95]. Quantitative analyses were performed as for RT-qPCR experiments using technical triplicate PCR samples. For ChIP-sequencing, a first ChIP series was performed using wild-type Col-*0*, *s2la-1*, *s2lb-2* and *s2la-1s2lb-2* seedlings, and a second series was performed using wild-type Col-*0* and *hub1-3* [49] plants, with an anti-H3K4me3 (Millipore #07-473) antibody before library preparation and Illumina sequencing. To ascertain that *hub1-3* plants used in these analyses displayed homozygous mutant alleles, seed batches from each corresponding stock were genotyped and "epigenotyped", the *HUB1* gene being reproducibly found among the few genes gaining H3K4me3 in *hub1-3* plants presumably because of T-DNA based ectopic transcription (Additional file 2 – Figure S13). Profiling of S2Lb-GFP and MYC-SDG2 was performed using anti-GFP (Life Technologies #11122) and anti-MYC (Ozyme #71D10) antibodies, respectively, and two crosslink steps. As recently described [96], samples were cross-linked first with 1.5 mM ethylene glycol bis(succinimidyl succinate) for 20 min and then with 1% formaldehyde for 10 min at room temperature. Cross-linking was stopped by adding 1.7 mL of 2 M glycine and incubating for 10 min. Libraries were prepared using 1 to 10ng of Input or IP DNA as described in the corresponding NCBI accession Super-Series GSE124319 datasets. TruSeq adapters were removed from sequenced short reads with trimmomatic v0.36 [92] using the following different parameters for each ChIP type: 1) for H3K4me3 in *s2l* mutants: “-phred33 LEADING:5 TRAILING:5”. Dataset-specific parameters were also used: “PE -- validatePairs ILLUMINACLIP:TruSeq2-PE.fa:2:30:10 MINLEN:20.”, 2) for H3K4me3 in *hub1.3* mutant: “SE ILLUMINACLIP:TruSeq3-SE.fa:2:30:10 MINLEN:30", 3) for S2Lb-GFP and MYC-SDG2: “PE-validatePairs ILLUMINACLIP:TruSeq3-PE.fa:2:30:10 MINLEN:20" Reads from all ChIP-Seq experiments were aligned to TAIR10 genome assembly with Bowtie2 v.2.3.3 [97] with “--very-sensitive” setting. Peaks were identified with MACS2 v2.1.1 [98] with the command MACS2 callpeak and different parameters for each ChIP-seq type: 1) for H3K4me3 in *s2lb* mutants: “-f BAMPE -- nomodel-q 0.01 -g 120e6 --bw 300”, 2) for H3K4me3 in *hub1-3* mutant: “macs2 callpeak -f BAM -q 0.01 -- bdg -g 120e6 --bw 300 --verbose 3 --nomodel -- extsize 200”, and 3) for S2Lb-GFP and MYC-SDG2: “-f BAMPE --nomodel -q 0.05 --bdg -g 120e6 --bw 300”. For S2Lb-GFP and MYC-SDG2 peak detection, the peaks found in the wild-type negative controls were used to clean up the peak lists of S2Lb and SDG2 profiles. H3K4me3 peaks obtained from two independent biological replicates were merged with bedtools v2.27.1 intersect [99]. Peaks from each experiment were annotated with Araport11 genes bedtools v2.27.1 intersect. Genes were considered marked by H3K4me3, S2Lb-GFP or MYC-SDG2 if overlapping for at least 150 bp with a relevant peak. To include nucleosomes in close proximity of the TSS, an upstream region of 250-bp was also considered for the overlap. Depth-normalized average values of read densities were computed over 10-bp non-overlapping genomic bins with Deeptools v3.1.0 bamCoverage [100] and used to draw the metagene plots and heatmaps with Deeptools computeMatrix, plotHeatmap and plotProfile. The normalized read densities of S2Lb-GFP, MYC-SDG2 and H3K4me3 in wild-type were also used to generate co-occurrence plots over the TSS of S2Lb-occupied genes using R 3.4.3 (www.R-project.org) and the package ggplot2 v3.1. 0 (www.github.com). Gene ontology analysis was performed using the GO-TermFinder software [101] via the Princeton GO-TermFinder interface (http://go.princeton.edu/cgi-bin/GOTermFinder). GO categories were filtered with the REVIGO platform [102].

## Authors’ contributions

All authors contributed reagents, materials or analysis tools. Protein analyses have been performed by ASF, MR, GZ, DS and AFD, except for MS analyses that were carried out by BL and supervised by DL. ASF performed DNA cloning, protein sequence analyses, microscopy, conducted RNA-seq experiments and all plant phenotyping analyses except dormancy assays that were conducted by EL and supervised by ChBa. RT-qPCR experiments were conducted by ASF and ClB. ChIPs were performed by ASF, ClB and MR. DL, AM and SKK prepared ChIP libraries and performed ChIP Illumina sequencing. RNA-seq library preparation and Illumina sequencing were performed at the Ecole normale superieure Genomic Platform (Paris, France). LC and ClB performed bioinformatics analyses. ASF, ClB, MB, ChBo and FB conceived the experiments. FB, MB and ChBo rose funding. ASF, ClB, ChBo and FB wrote the manuscript.

### Acknowledgements

We kindly thank Prof. Yuehui He (Shanghai Center for Plant Stress Biology, China) for providing *wdr5a-1* seeds, Prof. Xiaoyu Zhang (University of Georgia, USA) for providing *sdg2-3*/*SDG2::myc-SDG2* line and Dr Olivier Mathieu (CNRS, GreD, France) for providing the *Ds1-388-5* parental line. Authors are also much indebted to Alexandre Berr and Wen-Hui Shen (CNRS, Strasbourg, France) for critical reading of the manuscript and Vincent Colot (IBENS, Paris, France) for helpful discussions.

## Ethics approval and consent to participate

Not applicable.

## Consent for publication

All authors read and approved the manuscript.

## Availability of data and materials

All ChIP-seq and RNA-seq raw data generated in this work, except RNPII ChIP-seq datasets, are accessible through GEO Series accession number GSE124319. RNPII ChIP-seq datasets are available through GEO accession numbers GSM2028113 (ChIP) and GSM2028107 (Input). All ChIP-seq and RNA-seq gene lists are also given as supplementary tables 2 to 5. The mass spectrometry proteomics data are available via ProteomeXchange with identifier PXD012292. Other material and datasets will be made available on request.

## Additional file 2 - Supplementary figures

**Figure S1.**
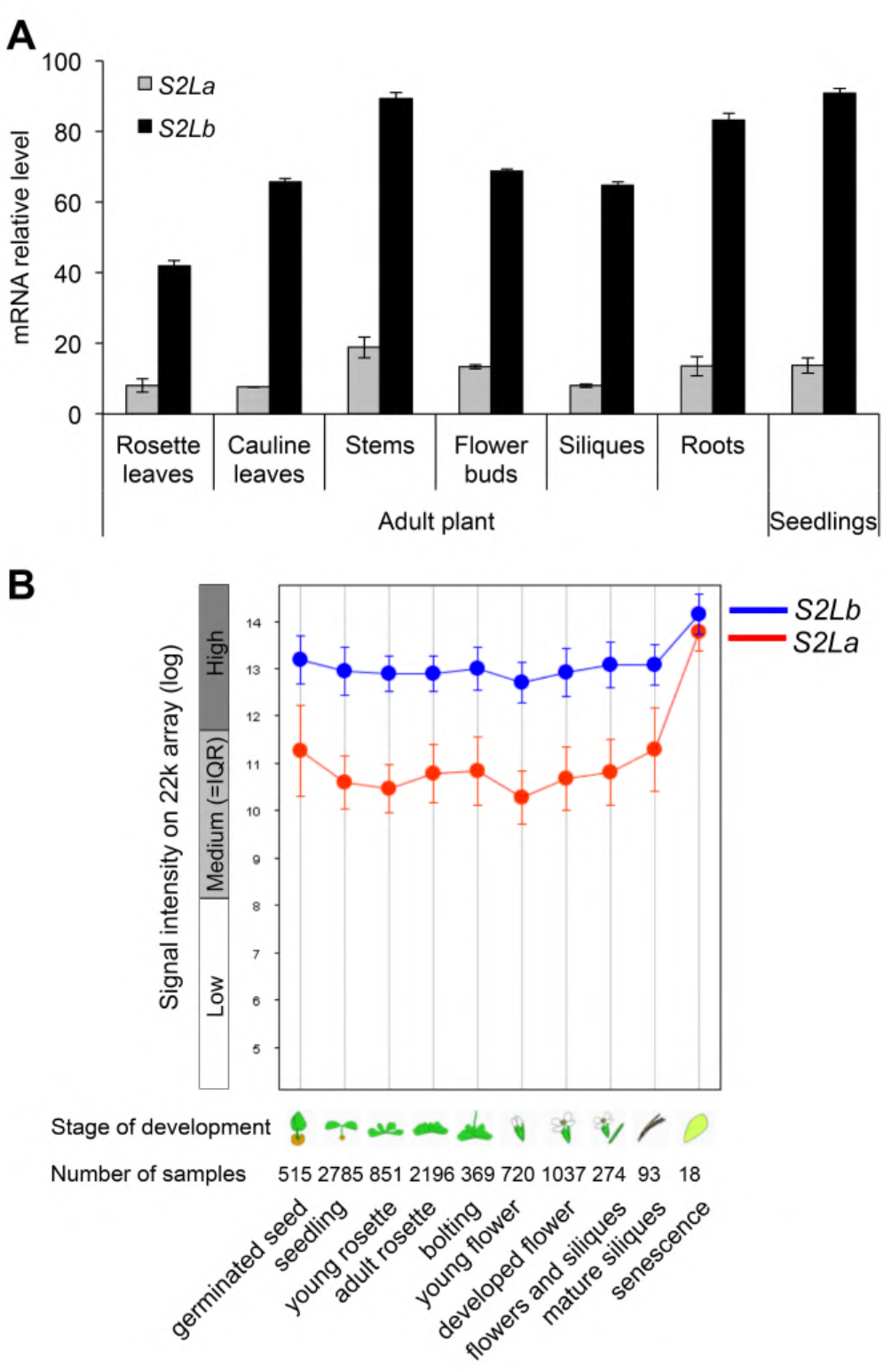
*S2La* and *S2Lb* mRNA levels in plant organs. **A)** Relative mRNA levels of *S2La* (grey bars) and *S2Lb* (black bars) in RT-qPCR analyses of different organs from from 10-day-old seedlings or wild-type adult plants (43 DAS). Error bars represent SD from two technical replicates. Transcript levels were normalized to the mean levels of *At2g36060*, *At4g29130* and *At5g13440* housekeeping genes. **B)** Genevestigator (Zimmermann et al., 2004) screenshot showing *S2La* and *S2Lb* gene expression at various developmental stages. Values are given as means +/-SD.

**Figure S2.**
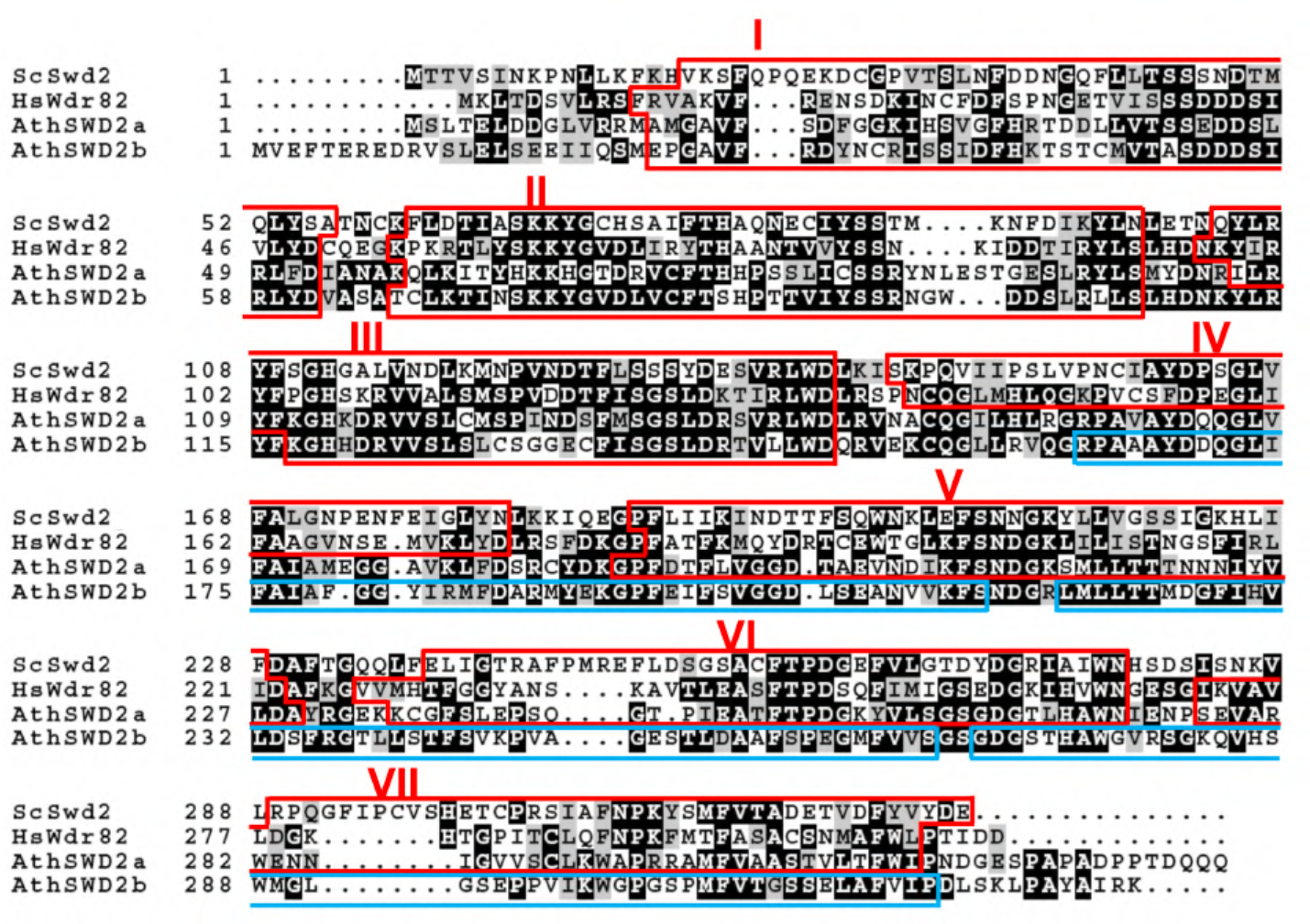
Amino-acids sequence alignment of full-length S2La and S2Lb with Swd2 (*S. cerevisiae*) and Wdr82 (*H. sapiens*) proteins. Numbers refer to amino-acids residues. Identical and similar residues are shaded in black and grey, respectively. WD40 repeats were defined according to WDSP predicting software (Wang et al., 2013). Conserved WD40 repeats are framed in red and numbered from I to VII. Divergent S2Lb WD40 repeats are framed in blue.

**Figure S3.**
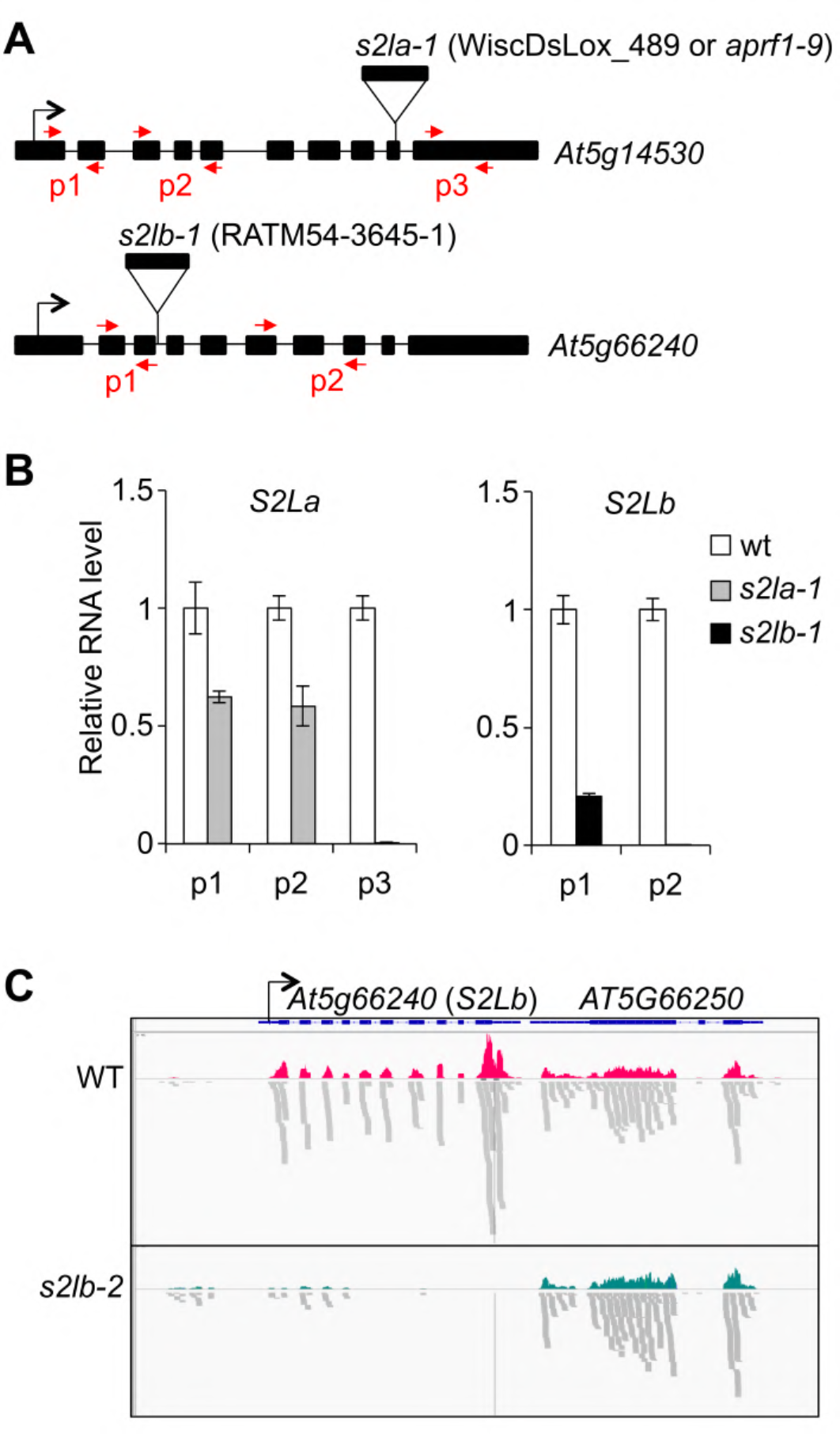
Expression of *S2La* and *S2Lb* genes in *s2la-1* and *s2lb-1* mutant plants. **A.** Schematic representation of the *S2La* and the *S2Lb* genes with the localization of the T-DNA insertions in the *s2la-1* and *s2lb-2* lines, respectively. Exons are represented by block boxes. **B.** RT-qPCR analysis of *S2La* and *S2Lb* transcript levels in adult *s2la-1* and *s2lb-1* mutant plants using the primers indicated (p1, p2, p3). RNA levels in the respective wild-type plants are arbitrarily set to 1. Primers positions are indicated in A (red arrows). Error bars represent SD from two technical replicates. Transcript levels were normalized to the mean levels of *At2g36060*, *At4g29130* and *At5g13440* housekeeping genes. **C.** Genome browser snapshot showing mRNA-seq read coverage (upper panels) and read positioning in wild-type and *s2lb-2* seedlings (lower panels).

**Figure S4.**
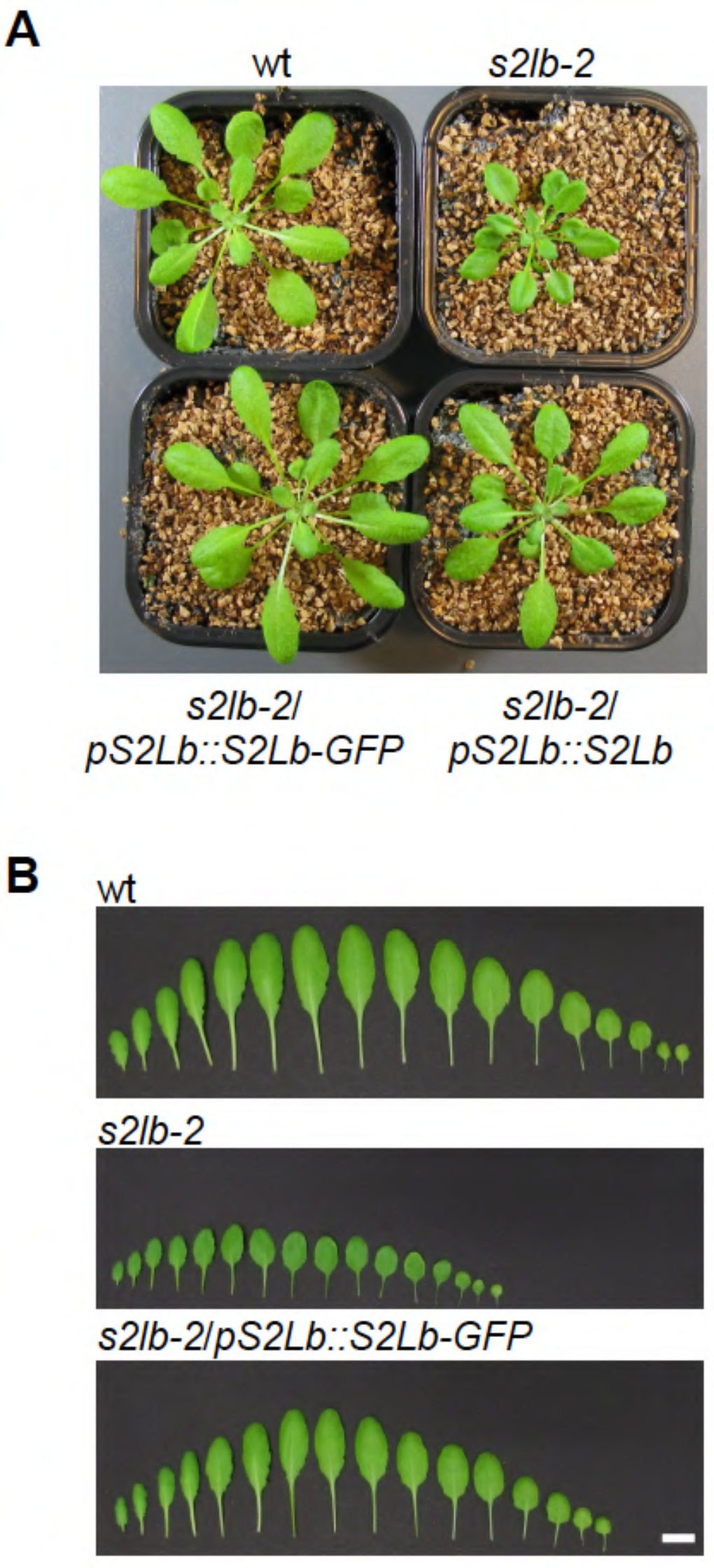
Complementation of *s2lb-2* morphological phenotypes by *pS2Lb::S2Lb* and *pS2Lb::S2Lb-GFP* constructs. Rosette **A)** and leaf **B)** phenotypes of 4-week-old plants grown under short day conditions. Scale bars, 1 cm.

**Figure S5.**
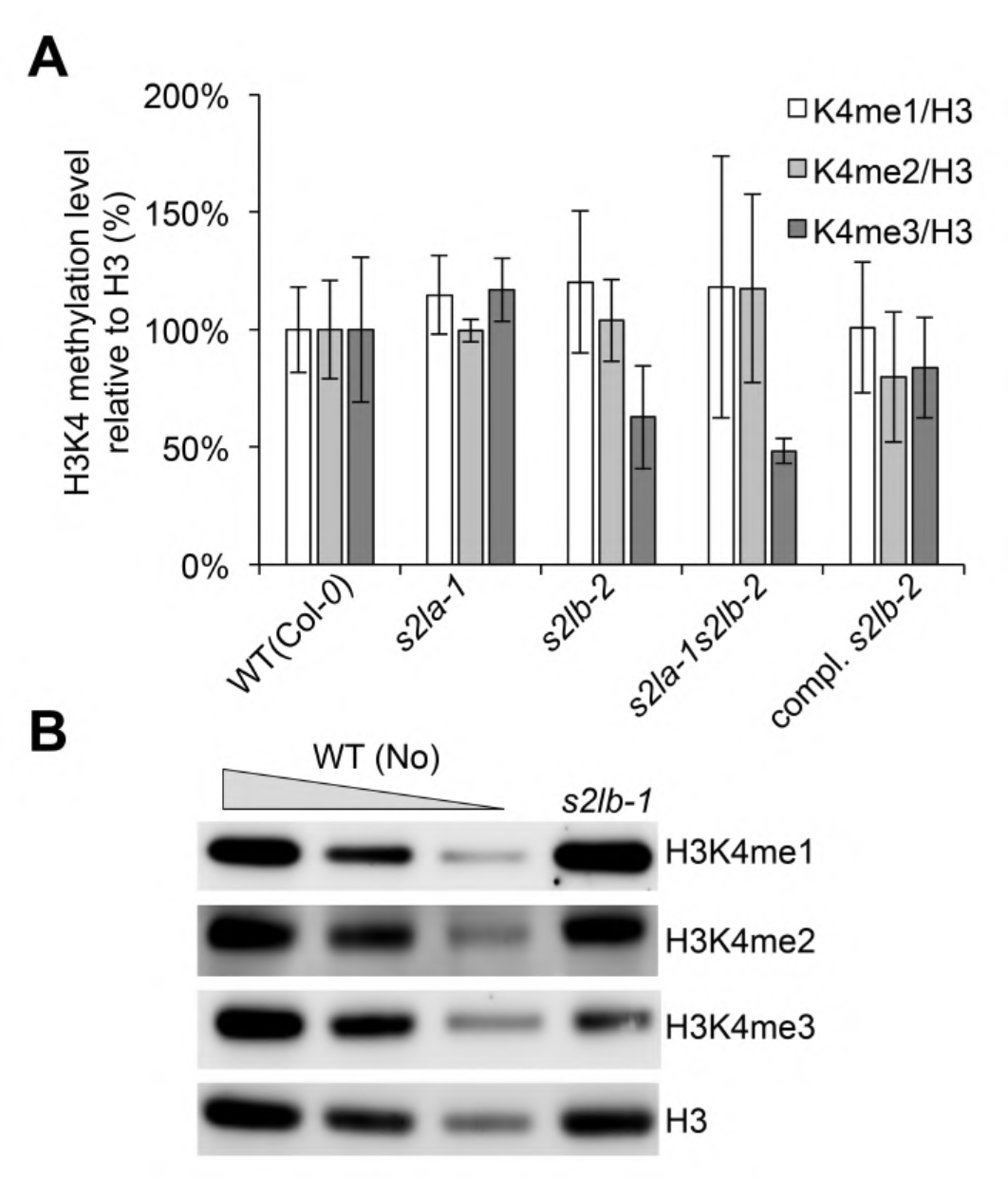
Decreased H3K4me3 level in *S2Lb* loss-of-function plants. **A)** Semi-quantitative analysis of H3K4 methylation signals in *S2L* loss-of-function plants. H3K4me1, H3K4me2 and H3K4me3 signal intensity relative to total histone H3 level were quantified using a luminescence imager. Levels in wild-type plants are arbitrarily set to 1. Error bars indicate SD from 3 independent biological replicates. **B.** Same as in (A) using antibodies recognizing H3K4me1, H3K4me2 and H3K4me3 and chromatin extracts from wild-type Nossen and *s2lb-1* plants. Histone H3 is used as loading control for H3K4me3 and H2Bub signals.

**Figure S6.**
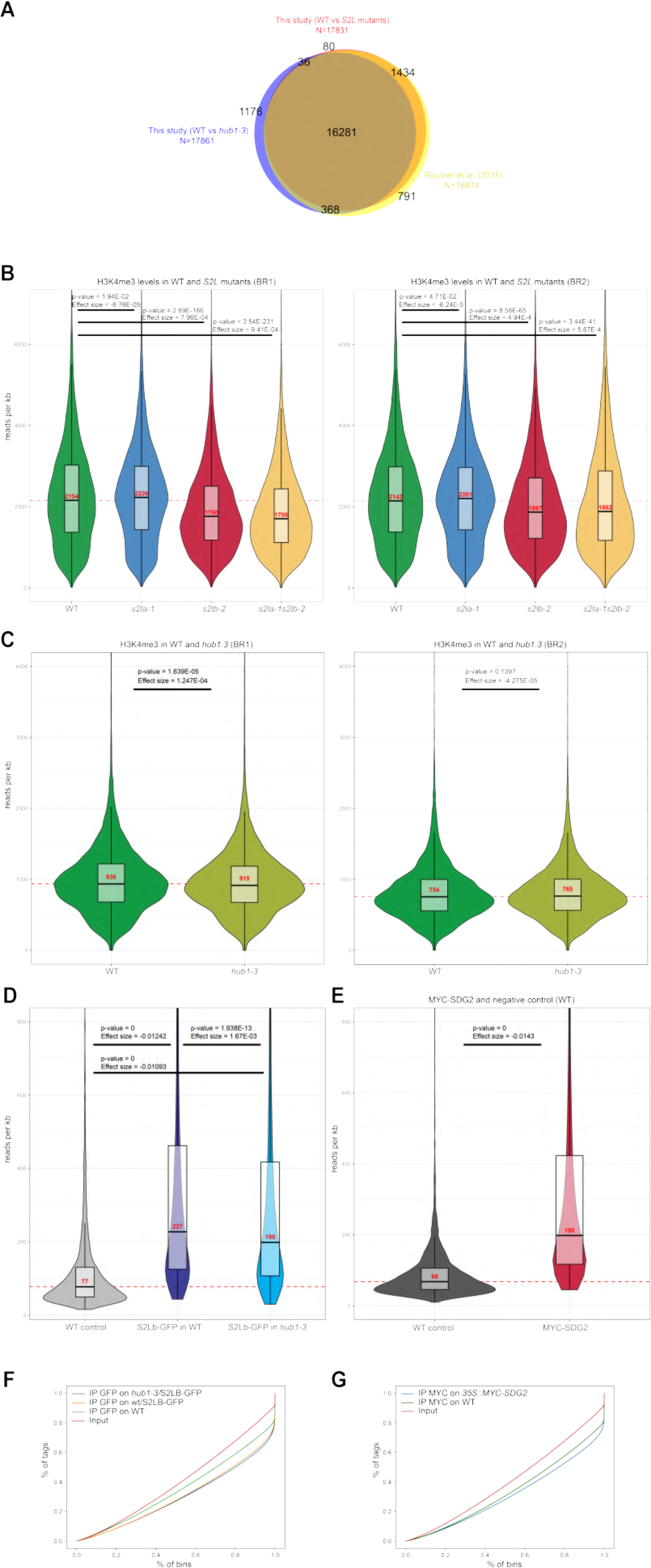
Validation and statistical analysis of ChIP-seq datasets. **A)** H3K4me3-marked genes identified in this study as compared to those in Roudier et *al.* (2011). The Venn diagram shows the overlap between H3K4me3-marked genes in this study (Additional file 3) and in Roudier et al (2011). **B)** Distribution of H3K4me3 short reads in wild-type and *S2L* mutant plants over genes overlapping with ChIP peaks identified in WT for the biological replicate 1 (BR1; left panel) and biological replicate 2 (BR2; right panel). Median read coverage is indicated in red. P-values for pairwise Mann–Whitney U test comparisons with the WT sample are given. The pValues indicate that *s2lb-2* and *s2la-1s2lb-2* mutants display significant lower levels than compared to WT, whereas *s2la-1* mutant plants have retained most of the mark. **C)** Same analysis than B) for *hub1-3* mutant plants. A measure of the effect size is also reported and indicates that the read density over marked genes is not different in mutant compared to wild-type plants. **D)** Distribution of short reads in wild-type plants (WT control), *WT/S2Lb::S2Lb-GFP* and *hub1-3/S2Lb::S2Lb-GFP* lines. For both transgenic lines, the IPs show a robust GFP signal enrichment as compared to the WT negative control, which is slightly weaker in *hub1-3* mutant plants. **E)** Same analysis than D) for MYC-SDG2 enrichment in *35S::MYC-SDG2* plants as compared to wild-type plants (WT control). The IP displays a strong enrichment in the transgenic line as compared to wild-type plants. **F)** Fingerprint analysis of S2Lb-GFP ChIP-seq. Cumulative percentage of tag in IP and Input samples over 500,000 randomly chosen 1000-bp genomic bins computer according to Diaz at al., 2012 (Stat. Appl. Genet. Mol. Biol., doi: 10.1515/1544-6115.1750). Reads are evenly distributed over all bins in Input and wild-type negative control samples, producing a linear cumulative distribution. Conversely, samples from the transgenic lines display a curve cumulative distribution, indicating that the highest-ranking bins (peaks, rightmost part of the curve) contain a larger fraction of short reads compared to the remaining bins (background). **G)** Same analysis than F) for MYC-SDG2 ChIP-seq.

**Figure S7.**
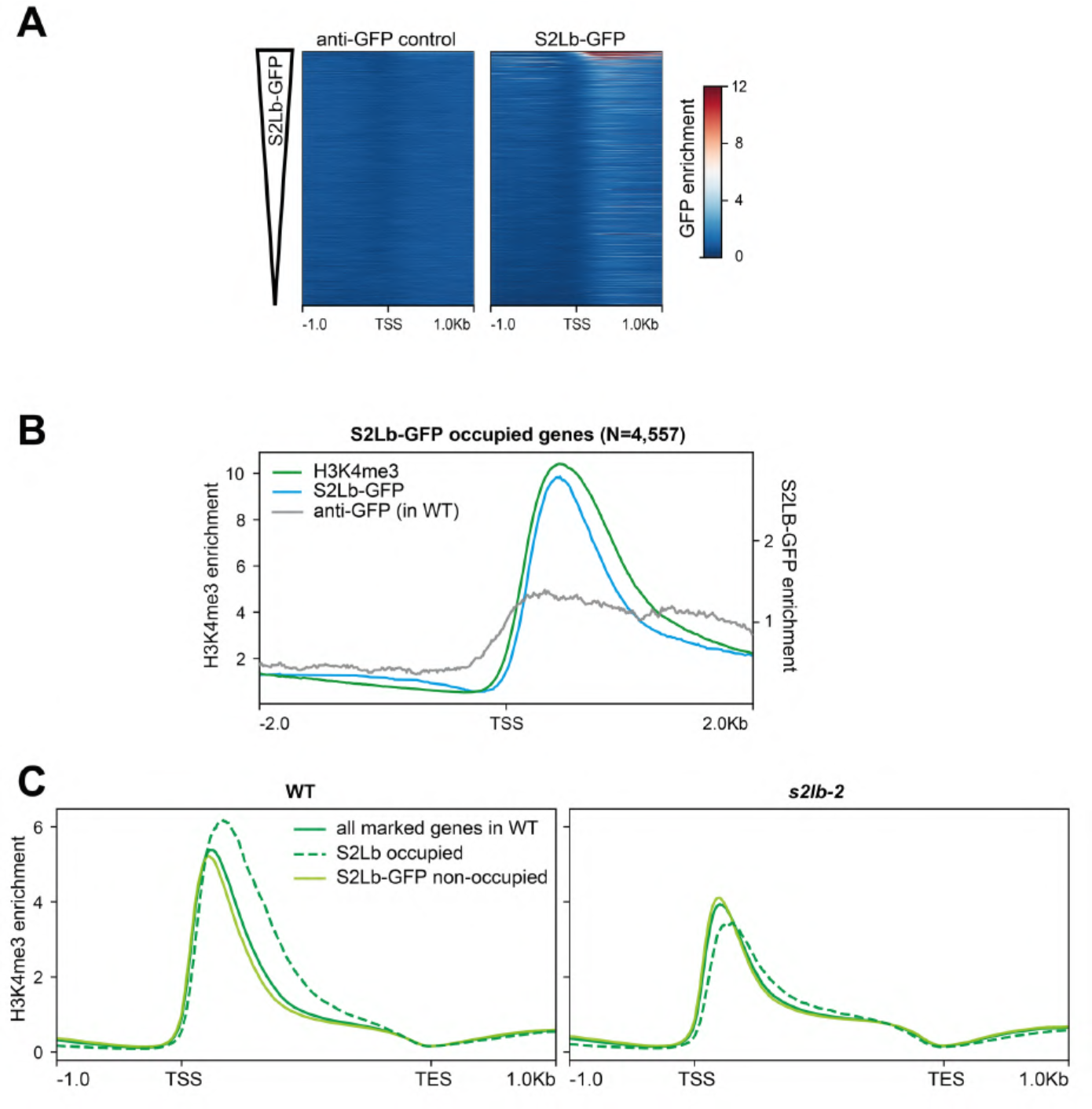
H3K4me3 over S2Lb-GFP targeted genes. **A) S2Lb** median enrichment over the genes marked by H3K4me3 in WT plants (N=17,831). Genes are equally ranked from top to bottom in each plant line according to median enrichment of S2Lb-GFP in the right panel. **B)**. Metaplot showing the median enrichment in H3K4me3 in WT plants (green line) or of S2LB-GFP (blue line) over the 4,557 genes targeted by H3K4me3. Background signal is shown by performing an anti-GFP ChIP-seq on wild-type plants grown simultaneously. **C)** Genes targeted by S2LB are globally highly marked by H3K4me3. Metaplots display H3K4me3 median enrichment in wild-type (left panel) or *s2lb-2* (right panel) over all H3K4me3-marked genes in WT seedlings (N=17,831; black line), and the ones occupied by S2Lb-GFP (N=4,557; yellow line) or marked by H3K4me3 but not occupied by S2Lb (N=13,455; blue line).

**Figure S8.**
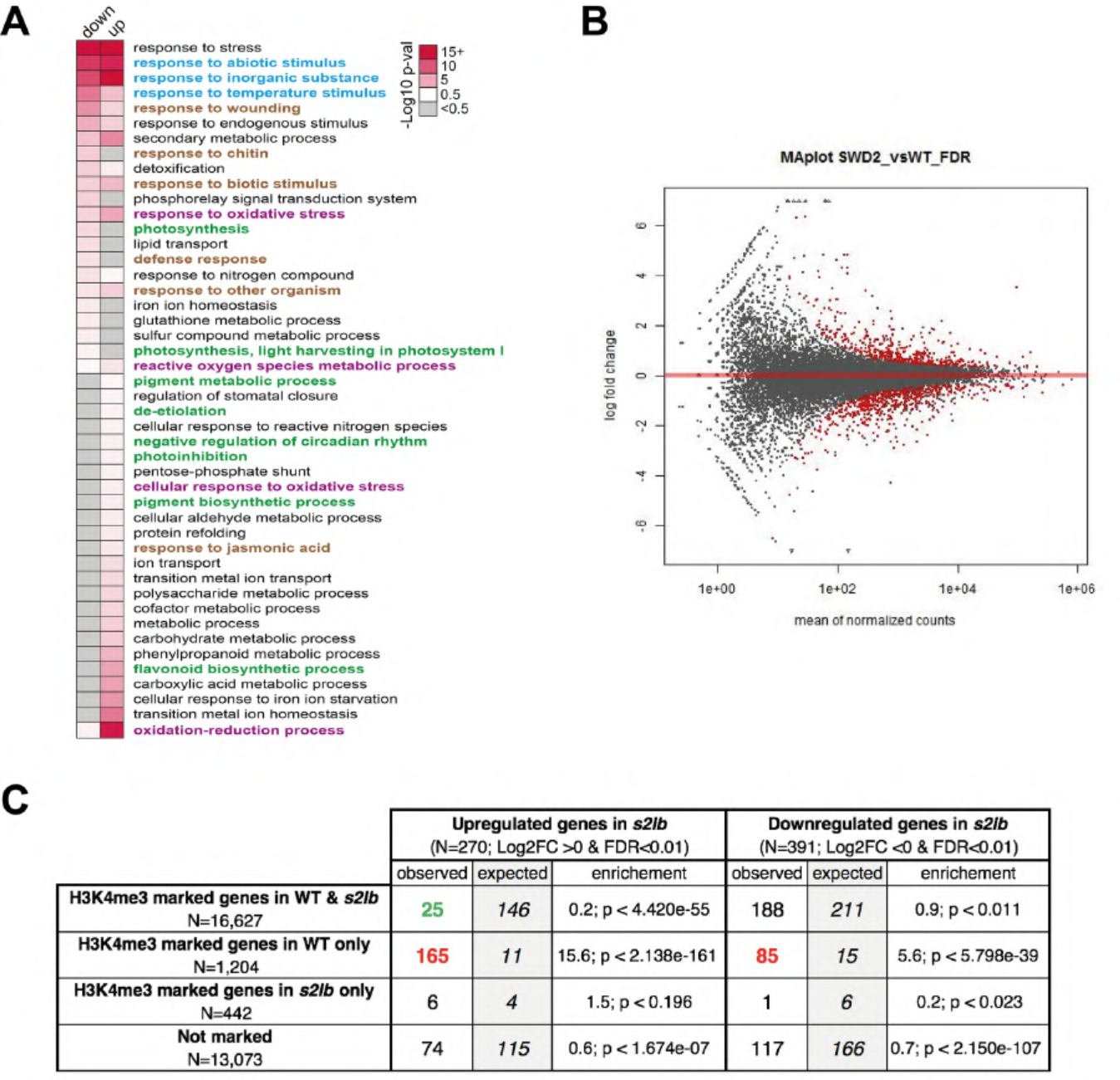
RNA-seq analysis of *s2lb-2* mutant seedlings. **A)** Heatmap showing the Gene Ontology (GO) enrichments (-Log10 p-Val < 0.5 in either up or downregulated genes) for DE genes in *s2lb-2* vs wild-type (FDR<0.01; Additional file 5). Grey color indicates -Log10 p-Val < 0.5. **B)** MA-plot showing the result of the DE-Seq2 analysis of the RNA-seq data comparing the transcriptome of wild-type and *s2lb-2* seedlings. 270 and 391 genes were found to be up and downregulated, respectively in the mutant plants compared to the wild-type (Additional file 5; FDR<0.01; red dots). **C)** Table showing the overlap between DE genes in *s2lb-2* vs WT (FDR<0.01; Additional file 5) and H3K4me3 marking in WT and s2lb. The enrichment p-value was calculated through the nemates.org website. Red color indicates a significant enrichment while green indicates a significant depletion.

**Figure S9.**
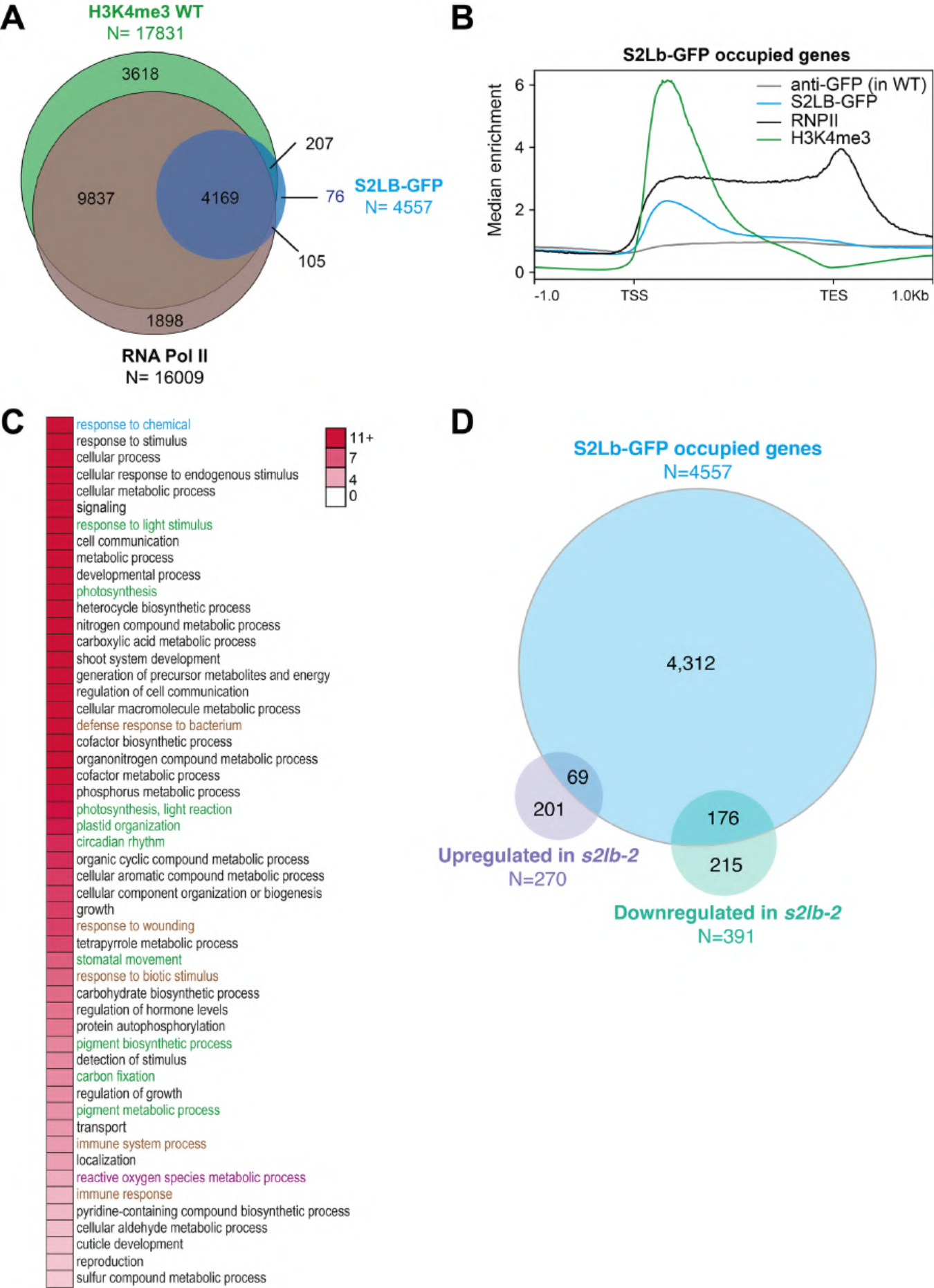
S2Lb-GFP expression and RNA-seq. **A)** Overlap between the genes occupied by S2Lb-GFP, H3K4me3 or by RNPII in WT seedlings. Overlap between H3K4me3 and RNPII, representation factor of 1.5 with p-Val<0.000e+00; overlap RNPII and S2Lb-GFP, representation factor of 1.8 with p-Val<0.000e+00; overlap between H3K4me2 and S2Lb-GFP, representation factor of 1.7 with p-Val<0.000e+00. **B)** Median enrichment of H3K4me3, S2Lb-GFP and RNPII over the 4,557 genes occupied by S2Lb-GFP. **C)** Heatmap showing GO terms enrichments (-Log10 p-value) for the 4,557 genes occupied by S2Lb-GFP. The category terms also found enriched for misregulated genes in *s2lb-2* are labeled with by the same color code. **D)** Overlap between the 4,557 genes targeted by S2Lb-GFP and the genes misregulated in *s2lb-2* seedlings (Additional file 5; FDR<0.01). The overlap is significantly enriched (upregulated genes, representation factor of 1.8 with p-Val<1.329e-06; downregulated genes, representation factor of 3.1 with p-Val<3.938e-48).

**Figure S10.**
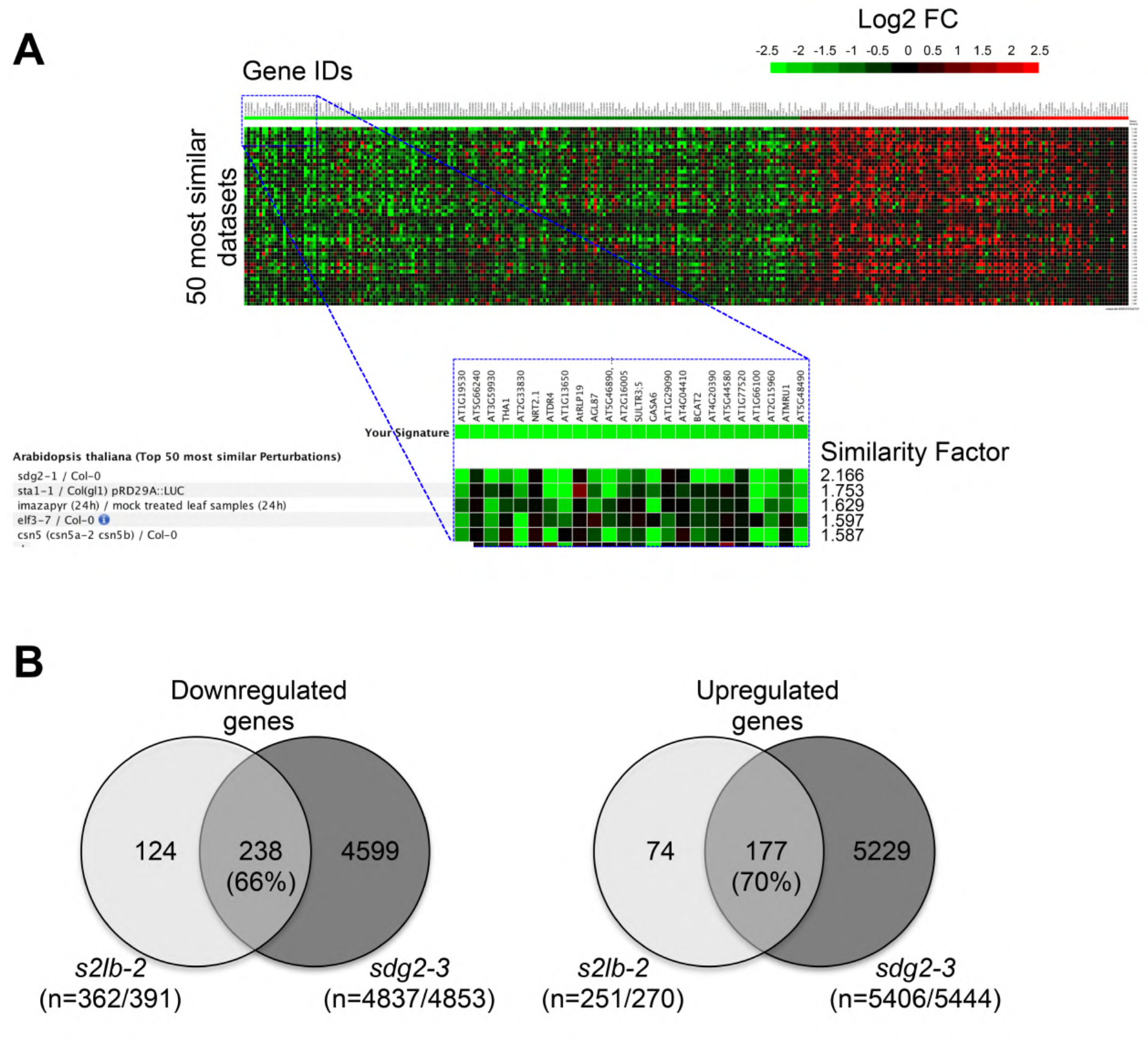
S2Lb and SDG2 co-regulate a large set of genes and associate within a high molecular weight complex. **A)** The Genevestigator Signature Tool was used to identify conditions that cause similar gene expression misregulation than the one found in *s2lb-2*. The expression values from genes misregulated in *s2lb-2* mutant seedlings (N=337, LogFC ≥ 1 & FDR ≤ 0.01) were compared to 3,282 other transcriptome experiments (ATH1 arrays) using the Pearson correlation to measure distances. The screenshot shows the 50 most similar datasets (from top to bottom). The inset shows an enlargement over the most downregulated genes in *s2lb-2* seedlings showing that the transcriptome profiling of the *sdg2-1* mutant transcriptome (AT-00435) from Guo et al. (2010) displays the strongest similarity. **B)** Venn-diagrams showing the comparison of down-or upregulated genes in *s2lb-2* (│LogFC│ > 0 and FDR ≤ 0.01) and *sdg2-3* (p-Val < 0.05) mutant seedlings. Only the gene sets for which data are available in different each experiment were kept for further analyses and both numbers are indicated in parentheses. The percentage of genes misregulated both in *s2lb-2* and *sdg2-3* datasets is indicated. The *sdg2-3* transcriptomic data are from (Guo *et al*, 2010).

**Figure S11.**
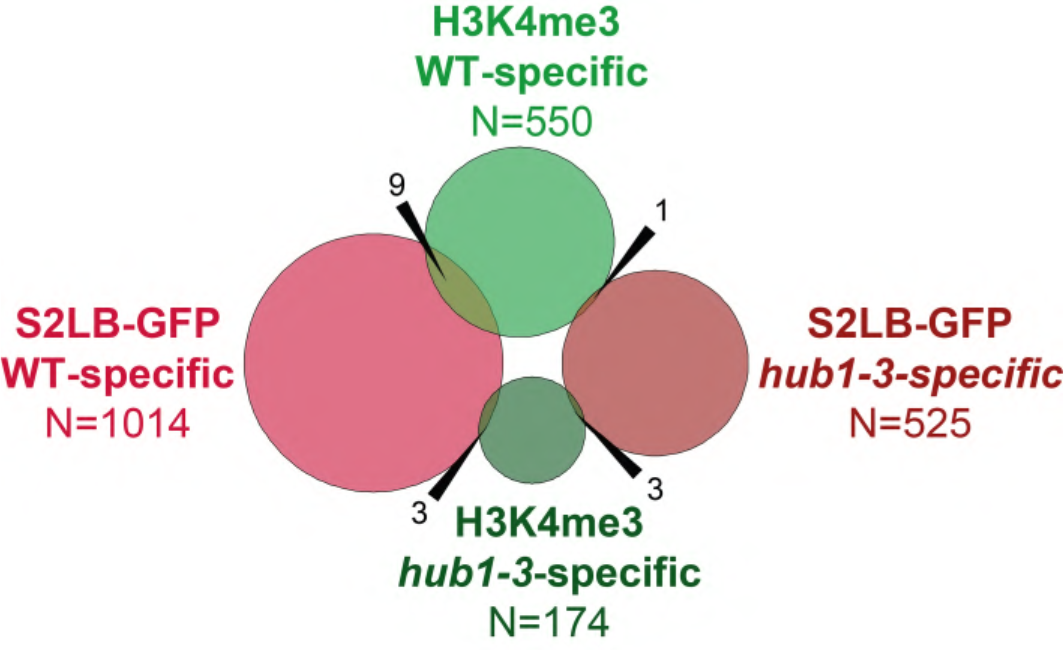
Detection of S2Lb-targeted genes losing a H3K4me3 peak in *hub1-3* seedlings. The Venn diagram shows the overlap between the lists of genes displaying specific marking by H3K4me3 or S2Lb in wild-type or in *hub1-3* seedlings. Only 9 genes appear to be subject to an H2B-H3 *trans*-histone crosstalk mediated by S2Lb, i.e. losing both H3K4me3 and S2Lb-GFP association in *hub1-3* plants. WT-specific: peaks identified in wild-type but not in *hub1-3* plants. *hub1-3*-specific: peaks identified in *hub1-3* but not in wild-type plants.

**Figure S12.**
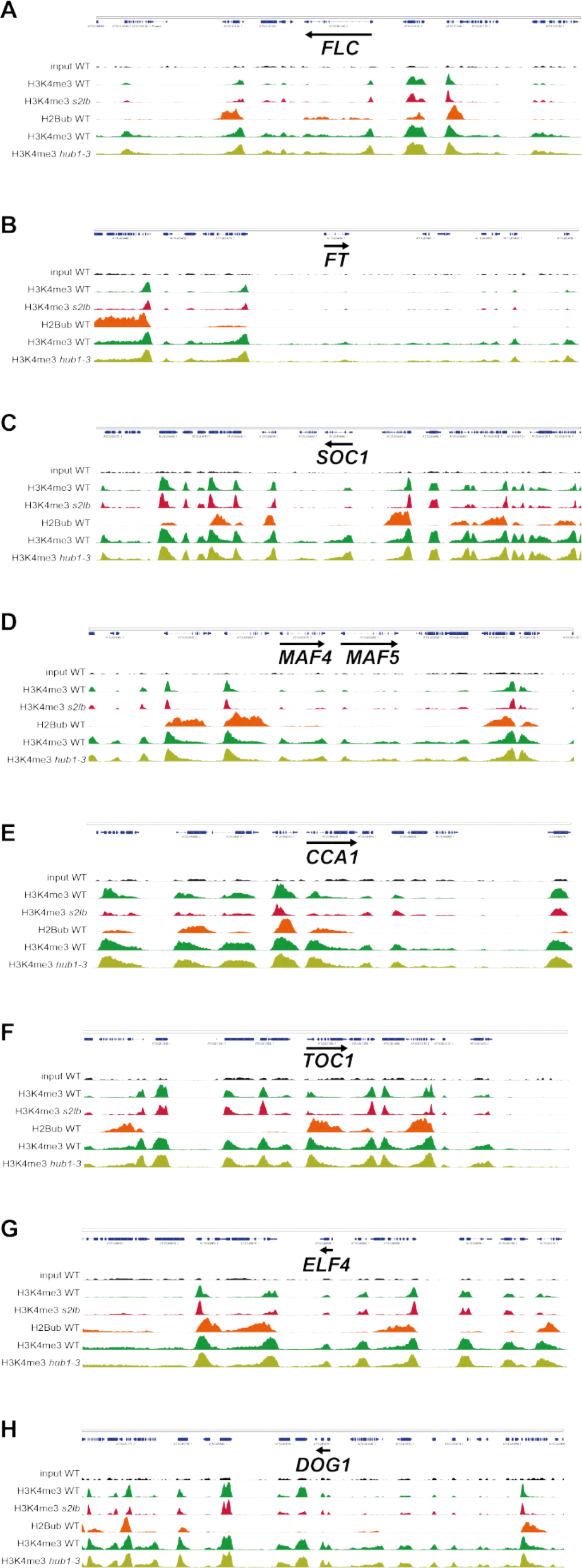
Genome browser snapshots of H3K4me3 and H2Bub profiles over representative genes in *s2lb-2* and *hub1-3* plants. H3K4me3 signals in different genetic backgrounds are equally scaled.

**Figure S13.**
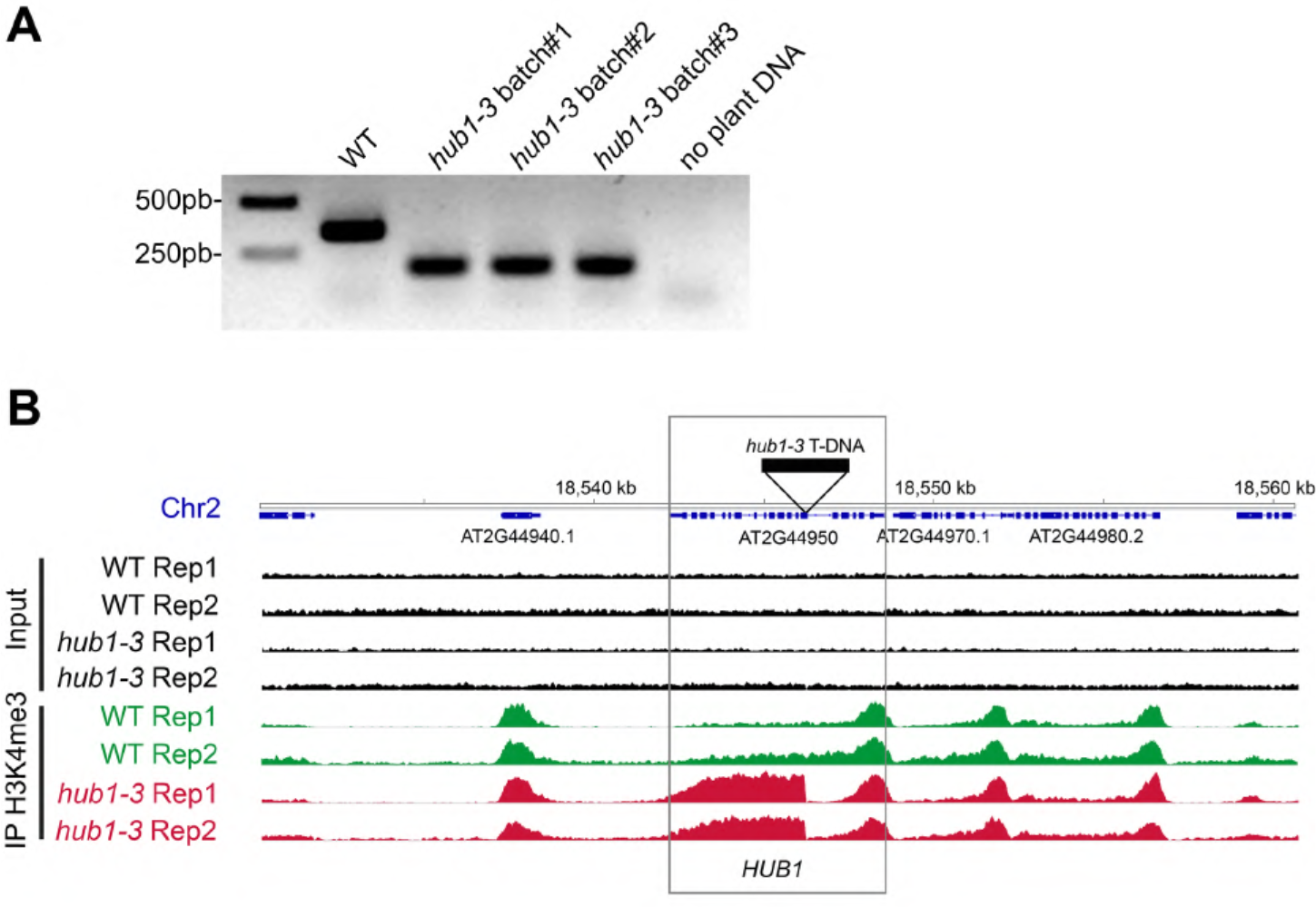
PCR genotyping and ChIP epigenotyping of the homozygous *hub1-3* allele in the seed batches used for ChIP-seq analyses in this study. **A)** PCR analysis of wild-type (WT) and *hub1-3* seed stocks used in H3K4me3 ChIP-seq experiments in Figure 6. DNA extraction and analysis were performed on a mix of 20 randomly picked seeds. **B)** H3K4me3 profile over the *HUB1* gene in the two independent biological replicates from the ChIP-seq analyses in Figure 6.

